# Proton export drives the Warburg Effect

**DOI:** 10.1101/2021.09.20.461019

**Authors:** Shonagh Russell, Liping Xu, Yoonseok Kam, Dominique Abrahams, Daniel Verduzco, Joseph Johnson, Tamir Epstein, Epifanio Ruiz, Mark C. Lloyd, Jonathan Wojtkowiak, Alex S. Lopez, Marilyn M. Bui, Robert J. Gillies, Pawel Swietach, Bryce Ordway

## Abstract

Aggressive cancers commonly ferment glucose to lactic acid at high rates, even in the presence of oxygen. This is known as aerobic glycolysis, or the “Warburg Effect”. It is widely assumed that this is a consequence of the upregulation of glycolytic enzymes. Oncogenic drivers can increase the expression of most proteins in the glycolytic pathway, including the terminal step of exporting H^+^ equivalents from the cytoplasm. Proton exporters maintain an alkaline cytoplasmic pH, which can enhance all glycolytic enzyme activities, even in the absence of oncogene-related expression changes. Based on this observation, we hypothesized that increased uptake and fermentative metabolism of glucose could be driven by the expulsion of H^+^ equivalents from the cell. To test this hypothesis, we stably transfected lowly-glycolytic MCF-7, U2-OS, and glycolytic HEK293 cells to express proton exporting systems: either PMA1 (yeast H^+^-ATPase) or CAIX (carbonic anhydrase 9). The expression of either exporter *in vitro* enhanced aerobic glycolysis as measured by glucose consumption, lactate production, and extracellular acidification rate. This resulted in an increased intracellular pH, and metabolomic analyses indicated that this was associated with an increased flux of all glycolytic enzymes upstream of pyruvate kinase. These cells also demonstrated increased migratory and invasive phenotypes *in vitro*, and these were recapitulated *in vivo* by more aggressive behavior, whereby the acid-producing cells formed higher grade tumors with higher rates of metastases. Neutralizing tumor acidity with oral buffers reduced the metastatic burden. Therefore, cancer cells with increased H^+^ export increase intracellular alkalization, even without oncogenic driver mutations, and this is sufficient to alter cancer metabolism towards a Warburg phenotype.

## Introduction

In 1924, Otto Warburg and colleagues demonstrated that cancer, even in the presence of oxygen, ferments glucose to lactic acid at high rates (Warburg et al., 1924), and this was contemporaneously confirmed by the Coris (Cori and Cori, 1925). Aerobic glycolysis, commonly termed the “Warburg Effect” in cancer (DeBerardninis and Chandel, 2020; Vaupel and Multhoff, 2020), is undeniably a hallmark of primary tumors and aggressive invasive disease (Hanahan and Weinberg, 2011). This preference of tumors for aerobic glycolysis is exploited in diagnostic PET imaging of fluorodeoxyglucose ( ^18^F-FDG) uptake (Kunkel et al., 2003). It is commonly believed that this increased fermentative glycolysis, and thus proton flux, is driven by oncogenes, such as RAS, MYC, HIF, and AKT (Kim et al., 2007; Wonsey et al., 2002), and that this augmented flux out-competes the ability of mitochondria to oxidize pyruvate, leading to the net production and export of lactic acid and reconversion of NADH to NAD^+^ to maintain redox balance. Hence, the Warburg Effect could solely be an epiphenomenon of oncogene activation, which is consistent with the observation that fermentation under aerobic conditions is energetically unfavorable and does not confer any clear evolutionary benefits (Vander Heiden et al., 2009).

We propose an alternative to this canonical view, building on principles of evolutionary dynamics (Gatenby and Gillies, 2004b). Aerobic glycolysis is such a commonly observed phenotype of aggressive cancers (Rizwan et al., 2013; Yu et al., 2015), we argue that it *MUST* confer some selective advantage for tumor growth (Damaghi et al., 2021). An inherent consequence of glycolysis is lactic acid production, and we propose that acid secretion *per se* renders cells more competitive, despite the energetic cost (Gatenby et al., 2006b; Gillies et al., 2008). Indeed, oncogenic drivers upregulate expression of proton exporting systems, e.g., sodium hydrogen antiporters, NHEs (Cardone et al., 2015; Cheng et al., 2019), carbonic anhydrase 9, CA-IX (Kopacek et al., 2005; Mahon et al., 2016; Takacova et al., 2010), and sodium-bicarbonate co-transporters, NBCs (Boedtkjer, 2019). Such activities will exacerbate the intra- to extracellular pH gradient, raising the intracellular pH (pHi) and acidifying the extracellular pH (pHe). These transporters have been associated with breast cancer aggressiveness (Beketic-Oreskovic et al., 2010). Additionally, glycolytic enzymes all exhibit significant pH-dependence (Persi et al., 2018), and thus maintenance of an alkaline pHi would directly promote increased glycolytic flux.

Simultaneously, lowering pHe provides invading cancer cells competitive benefits that can enhance colonization, invasion, and metastasis. These benefits include extracellular matrix remodeling via release and activation of proteases to increase invasion (Kato et al., 2005; Stock and Schwab, 2009; Webb et al., 2011b), inhibition of immune surveillance (Brand et al., 2016), promotion of an epithelial-to-mesenchymal transition (EMT) (Nieto et al., 2016; Pastushenko et al., 2018; Puisieux et al., 2014; Riemann et al., 2019), and anchorage-independent growth (Damaghi et al., 2015; Jin et al., 2018; Paoli et al., 2013; Peppicelli et al., 2019). This is further supported by the observation that neutralization of pHe acidity can inhibit invasion and metastases (Estrella et al., 2013; Robey et al., 2009).

The current study investigates whether aerobic glycolysis can be driven by proton-export, and further investigates the impact of this on cancer aggressiveness. We demonstrate that over-expression of proton exporters is sufficient to increase aerobic glycolysis, through enhanced glucose uptake and lactate production. We further observed that proton export increased intracellular pH and increased metabolic flux at most steps in glycolysis. Finally, we observed *in vivo* that these proton-exporting cell lines were more aggressive, generating higher-grade tumors and increased metastases. There is a known association between acid production, aerobic glycolysis, and metastatic potential. Further, experimental metastases can be inhibited with acid-neutralizing buffers. The current work adds to this literature by demonstrating that acid production *per se* can be sufficient to drive the Warburg Effect and promote metastasis.

## RESULTS

### Overexpression of CA-IX in cancer cells increases glycolytic metabolism

CA-IX hydrates extracellular CO_2_ to H^+^ + HCO_3_^-^. This facilitates CO_2_ diffusion away from the cell reducing pH gradients across tissues. The bicarbonate generated from CO_2_ hydration, a reaction that occurs either spontaneously or is sped up by carbonic anhydrases, can then reenter the cell via Na^+^ + HCO_3_^-^ co-transporter (Boedtkjer, 2019; Svastova et al., 2012).

Numerous studies in, e.g., breast, ovarian (Choschzick et al., 2011), and astrocytoma (Nordfors et al., 2013) cancers have shown that CA-IX expression correlates with poor prognosis and reduced survival. Figure 1A shows the overall survival of ER+ breast cancer patients with low and high (median cutoff) CA-IX expression was 143 and 69.4 months, respectively, p=9.32 e-7. Although the sample size was smaller, a similar pattern was seen for metastasis-free survival in patients with low and high CA-IX expression, 130 vs. 50 months, respectively, p=0.0012. Metastasis-free survival is also reduced in other cancers with high CA-IX expression, including cervical and colorectal (van Kuijk et al., 2016). CA-IX’s role in outcome makes it a clinically significant target warranting further investigation.

**Figure 1:**
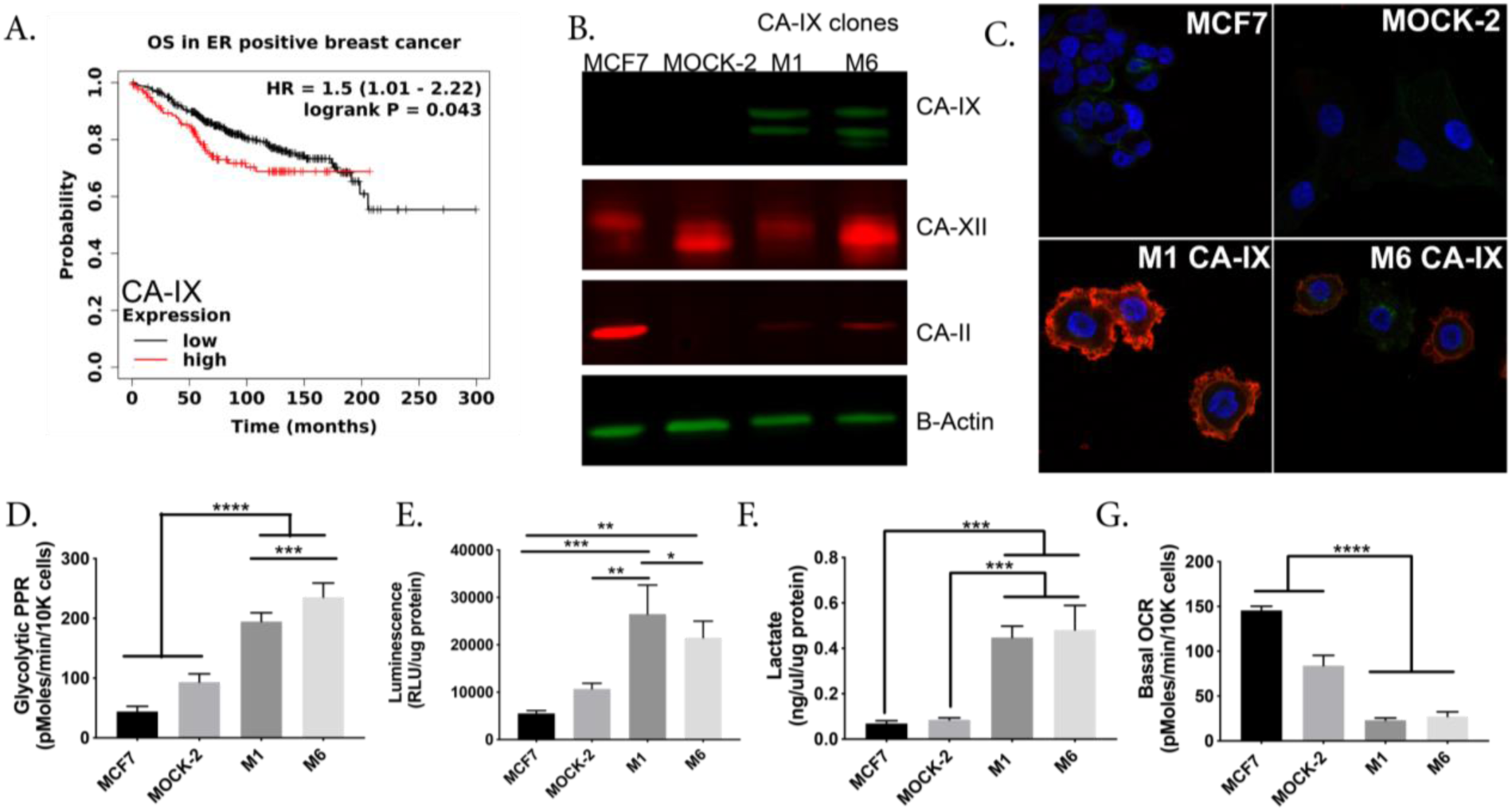
Over-expression of CA-IX in MCF-7 breast cancer cells increases glycolytic metabolism in vitro. **A:** Overall survival Kaplan-Meier Curve in ER-positive breast cancer comparing low and high CA9 gene expression n=1402 (kmplot.com). Statistical analysis using Log rank P test p=0.0024. **B:** Immunoblotting of protein lysates from MCF-7 cells transfected with empty vector (MOCK-2) or Ca9 vector (M1 and M6). Proteins from total cell extracts were immunoblotted for CA-IX, CA-II, CA-XII, and B-actin (loading control). **C**: Representative immunocytochemistry images of CA-IX protein expression in MCF-7, MOCK-2, and CA-IX clones M1 and M6. CA-IX clones (M1 & M6) exhibit CA-IX membrane staining, whereas MOCK-2 and parental MCF-7 cells do not. DAPI nuclear stain (blue), Wheat germ agglutinin membrane stain (green), and CA-IX stain (red). **D:** Glycolysis associated proton production rate (PPR) using the Seahorse extracellular flux analyzer, measured post glucose injection. Data are shown as mean ± SD, N=8 biological replicates per group, statistical analysis using ordinary one-way ANOVA. **E:** Glucose uptake of cells in each group over 24hr, measured as luminescence generated using Glucose Uptake-Glo assay (Promega). N=3, statistical analysis using ordinary one-way ANOVA. **F:** Lactate measured in extracellular media after 24hr using Sigma kit. N=3 biological replicated per group, statistical analysis using ordinary one-way ANOVA. **G:** Basal oxygen consumption rate (OCR) measured using the Seahorse extracellular flux analyzer in 5.8mM glucose concentration. Data are shown as mean ± SD, N=8 biological replicates per group, statistical analysis using ordinary one-way ANOVA. *p<0.05, **p<0.01, ***p<0.001, ****p<0.0001

To test the hypothesis that proton export can drive aerobic glycolysis, we established models which over-expressed proton exporters. We transfected MCF-7 cells with a CA-IX expression vector and isolated two individual clones (M1 and M6) and confirmed CA-IX protein expression (Fig. 1B). MCF-7 cells do not express CA-IX under normoxic conditions, but do express other carbonic anhydrases, CA-II and CA-XII (Fig. 1B). CA-IX is distinct among exofacial CA’s (CA-IV, CA-XII), as it contains a proteoglycan domain, which enables it to maintain enzymatic activity at lower pHe (Li et al., 2011). In tissues, CA-IX can function as a “pH-stat”, which tumors hijack to maintain an acidic pHe (Lee et al., 2018). We also transfected MCF-7 cells with an empty pcmv6 vector hereafter referred to as MOCK-2. Additionally, we confirmed by ICC that CA-IX, an exofacial membrane-bound protein, was expressed on the plasma membrane in both CA-IX clones (Fig. 1C).

To test our hypothesis that proton export can drive aerobic glycolysis, we interrogated the metabolism of our CA-IX expressing clones using a Seahorse XFe96 Extracellular Flux (XF) Analyzer, enzymatic, and radiochemical assays to assess both glycolytic and mitochondrial metabolism. Specifically, the Seahorse glycolytic stress tests (GST) showed that both CA-IX clones exhibited higher proton production rates (PPR) upon glucose stimulation compared to MOCK-2 or parental clones (Fig. 1D). Using glucose, and [^3^H]-2-deoxyglucose (2DG) uptake assays (Fig. 1E **&** Supplementary Fig. 1), and lactate production rate assays (Fig. 1F **&** Supplementary Fig. 2), we further confirmed CA-IX expressing clones had increased glycolysis in normoxic conditions. Mitochondrial metabolism in these clones, as measured by Seahorse mitochondrial stress test (MST), exhibited a decreased reliance on oxidative phosphorylation. In the CA-IX clones, both basal oxygen consumption rates (Fig. 1G) and reduced ATP-linked oxygen consumption rates (Supplementary Fig. 3) were decreased compared to MOCK-2 or parental cells. CA-IX expressing clones also had hyperpolarized mitochondria (Supplementary Fig. 4). Therefore, CA-IX expression did not globally upregulate all ATP turnover, but upregulated aerobic glycolysis and limited reliance on oxidative phosphorylation.

To investigate whether these metabolic alterations were specific to the MCF-7 cells, we tested other cell lines. CA-IX was over-expressed in U2-OS osteosarcoma human cells and HEK 293 human embryonic kidney cells. CA-IX expression upregulated glycolysis in both cell lines, as seen by increased aerobic lactate production (Supplementary Figs. 5 & 6). Even in HEK 293 cells, which have higher basal glycolysis than the other cell lines tested, over-expression of the proton exporting CA-IX still enhanced their rate of aerobic glycolysis.

To delineate which steps in glycolysis were being impacted by acid export, we analyzed the intracellular metabolites using mass spectrometry and a library of known metabolites (see methods). Principal component analysis (PCA) showed a statistically significant separation of MOCK-2 and parental MCF-7 from the CA-IX clones (Fig. 2A). Notably, PCA also showed that the parental and MOCK-2 cells had distinct metabolic profiles, but neither overlapped with the CA-IX clones. Consistent with this, heatmap visualization of the top 50 most significantly altered metabolites showed that the two CA-IX clones exhibited similar metabolic profiles (Fig. 2B) but were considerably different from the MOCK-2 and parental clones. MOCK-2 cells grow more rapidly *in vitro* compared to the parental MCF-7 or the CA-IX clones which is likely why they exhibit a different metabolic profile from the parental line (Supplementary Fig. 7). We note later, however, that this difference in growth rate was not maintained *in vivo* when grown as primary tumors (Fig.4A). Out of all metabolites assessed, the glycolytic intermediates were consistently altered in the CA-IX clones compared to parental or MOCK-2.

**Figure 2:**
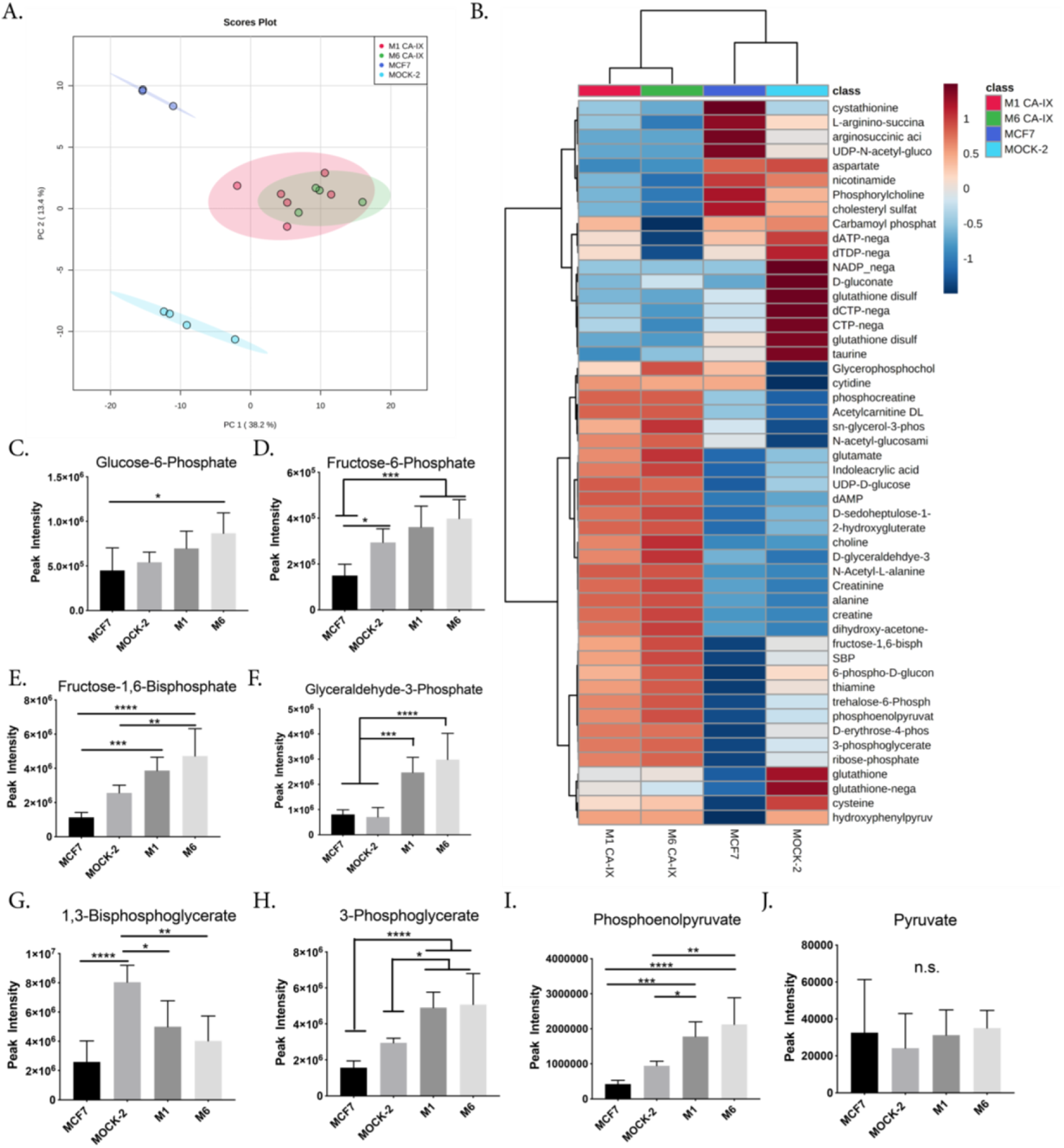
Unlabeled metabolic profiling of CA-IX expressing MCF-7 cells intracellular metabolites and analysis of the glycolytic metabolites. **A:** Principal component analysis of intracellular metabolites in CA-IX expressing, MOCK-2, and Parental MCF-7 cells. **B:** Heatmap (hierarchical clustering) of the fifty most significant fold changes of intracellular metabolites between CA-IX expressing clones, MOCK-2, and Parental cells. Numerous glycolytic intermediates were significantly higher in CA-IX expressing cells compared to MOCK-2 and Parental. **C-F:** Average peak intensity of each glycolytic intermediate in CA-IX expressing, MOCK-2 and Parental MCF-7 Cells. N=5-6 biological replicates per group, statistical analysis using ordinary one-way ANOVA. **G-J:** Average peak intensity of each glycolytic intermediate in CA-IX expressing, MOCK-2 and Parental MCF-7 Cells. N=5-6 biological replicates per group, statistical analysis using ordinary one-way ANOVA. *p<0.05, **p<0.01, ***p<0.001, ****p<0.0001

The CA-IX clones exhibited increased levels of all glycolytic intermediates upstream of pyruvate kinase, PK (Fig. 2C-J), which catalyzes the penultimate step of glycolysis: the conversion of phosphoenolpyruvate (PEP) + ADP → pyruvate +ATP. Thus, it appears that the activities of the upstream enzymes have increased, leading PK to now become rate-limiting for glycolytic flux in the CA-IX expressing cells (Fig. 2J). As the CA-IX cells have higher glycolytic flux (Fig. 1D-F), the most straightforward interpretation is that CA-IX expression de-inhibited all of the enzymatic steps upstream of PK. Overall, CA-IX expression enhances glycolytic intermediates’ flux, resulting in enhanced lactate and acid production.

### CA-IX over-expression increases pHi

Our metabolomics studies suggest multiple glycolytic enzymes were impacted, we therefore hypothesized that CA-IX expression might raise the intracellular pH of cells, which could pleiotropically increase glycolytic enzyme rates. Most glycolytic enzymes have ionizable residues that can alter their enzyme activity. Recently, these residues have been characterized for all glycolytic enzymes through homology modelling, which predicted glycolytic enzyme activities generally increase with pHi above neutral (Persi et al., 2018). We tested the PPR in MCF-7 parental cells after altering pHi in a range of pHe media 6.6-7.4, expecting that the PPR rate would increase with increasing pHi. Cells were incubated with either a chloride-containing solution or an iso-osmotic low-chloride formulation which replaced chloride salts with gluconate equivalents. Low-chloride solutions alter the driving force for Cl^-^/HCO_3_^-^ exchangers, effectively loading cells with HCO_3_^-^ ions and raising pHi at the same pHe (Sasaki and Yoshiyama, 1988; Wu et al., 2020). At all values of pHe from 6.6-7.2, cells in gluconate media exhibited significantly higher glycolytic rates compared to those in chloride-containing media (Fig. 3A **&** Supplementary Fig. 8), indicating that the increased glycolytic rate is strongly dependant on pHi.

**Figure 3:**
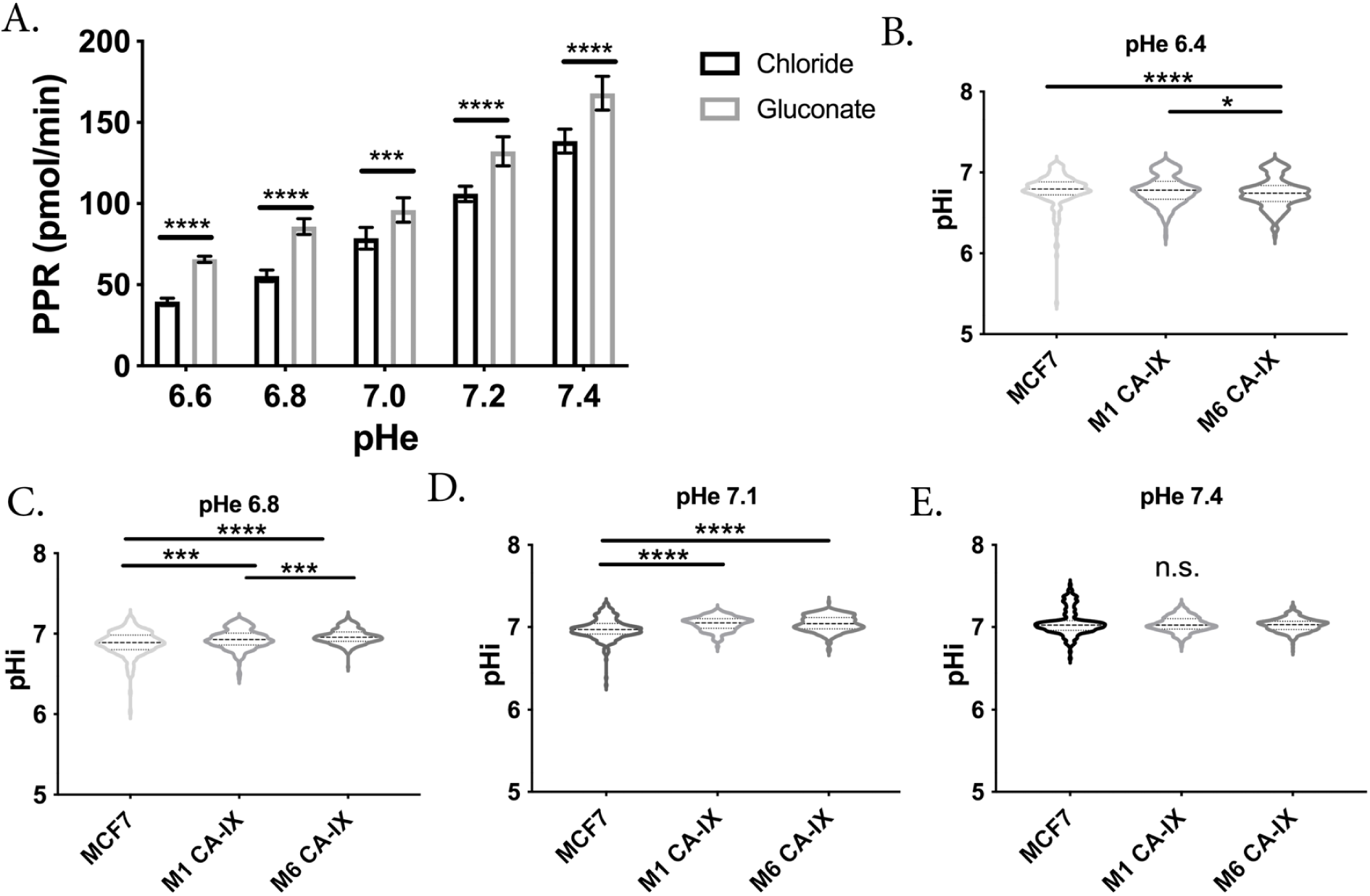
Increased intracellular pH enhances glycolysis in breast cancer cells. **A:** Increased intracellular pH, using gluconate substituted media, increases the glycolytic rate in MCF7 cells irrespective of pHe. N=8 per group, statistical analysis using Welch’s T-test; **B-E:** Intracellular pH as measured using cSNARF1 in varying extracellular pH, n=158-438 cells analyzed per group, statistical analysis using Kruskal-Wallis test. *p<0.05, ***p<0.001, ****p<0.0001

CA-IX overexpression has been shown to increase pHi in other systems (Morgan et al., 2007), leading us to probe pHi in our CA-IX expressing cells. We used a pHi reporter dye, cSNARF1 to measure the effects of CA-IX expression on intracellular pH (pHi). A cSNARF1 calibration curve was generated using nigericin/K^+^ buffers (Supplementary Fig. 9). We then loaded MCF7 parental or the CA-IX expressing clones with cSNARF1 and used fluorescence imaging to measure pHi. MOCK-2 cells were unsuccessfully loaded with cSNARF1, possibly due to reduction or loss of esterase activity from the integration of the MOCK-2 plasmid into the MCF-7 DNA; therefore, they are not included in these analyses. We equilibrated the cells in media at pHe 6.6, 6.8, 7.1, and 7.4 and co-loaded them with cSNARF1 and nuclear dye Hoechst 33342. The inclusion of a nuclear dye allowed post-processing to mask the nucleus, ensuring measurement of cytoplasmic pH only. At the intermediate pHe values of 6.8 and 7.1, the CA-IX expressing clones had significantly higher pHi compared to parental (Fig. 3C, D **&** Supplementary Table 1). The enzymatic activity of CA-IX is optimal at pH 6.8 and drops precipitously as the pH is lowered to 6.0 (Li et al., 2011)(McIntyre et al., 2012). Consistent with this, at an acidic pHe of 6.4, we observed that the MCF7 parental cells had a higher intracellular pH compared to CA-IX clones (Fig. 3B **&** Supplementary Table 1), suggesting CA-IX enzymatic function was strongly reduced resulting in reduced CO_2_ venting. There were no significant differences in pHi at pHe 7.4 (Fig. 3E **&** Supplementary Table 1). These data show that CA-IX expression can raise the pHi at intermediate pHe values and that thus increasing pHi likely results in increased glycolytic flux by globally enhancing glycolytic enzyme activity.

### CA-IX expression increases metastasis

We then investigated the effect of CA-IX expression on migration, invasion, and metastasis, as there are reported correlations between increased glycolytic flux, reduced extracellular pH, and metastasis (Birchmeier et al., 2003; Estrella et al., 2013; Hiraga et al., 2013; Kato et al., 2004). Additionally, CA-IX has been associated with increased invasion and metastasis in several systems (Swayampakula et al., 2017; Ward et al., 2015).

*In vitro*, we utilized scratch assays and gel escape to measure migration and invasion in the CA-IX clones. Scratch assays showed that clone M1 had increased migratory ability (Supplementary Fig. 10), closing the wound significantly more rapidly than controls. However, this was not observed in the M6 CA-IX clone. In the gel escape assay, expansion out of the gel is due to a combination of proliferation, invasion, and migration. Both CA-IX clones invaded the area surrounding the gel drop substantially quicker than the parental MCF-7 clone (Supplementary Fig. 11). Compared to the MOCK-2 cells, however, only the M6 clone was significantly more invasive. The apparent increased invasion rate by MOCK-2 compared to parental is likely due to their increased proliferation rate as correcting for proliferation eliminates the difference between MOCK-2 and parental (Supplementary Fig. 7). It is important to note that these assays were carried out at neutral pH and that the invasive behavior might be further enhanced by low pH, as we have shown previously for melanoma cells (Moellering et al., 2008). Together, these studies suggest that CA-IX expression can enhance cell motility, increasing migration and local invasion.

Furthermore, aggressive cancers with stem-like properties can resist anoikis, which can be measured *in vitro* by cells’ ability to form spheroids independent of attachment to a basement membrane. Utilizing the hanging droplet technique, we observed that CA-IX expression enabled robust, compact spheroid formation, compared to the MOCK-2 and parental MCF-7 clones, which could not (Supplementary Fig. 12). This spheroid forming ability suggests CA-IX not only enhances cell:cell adhesion but also suggests that increased proton export can contribute to anoikis resistance when detached from the basement membrane.

Although the phenotype of our proton exporting CA-IX clones appeared to be more aggressive, this could be an *in vitro* only phenomenon. We thus investigated the clones *in vivo.* We studied the effect of CA-IX expression on primary tumor growth, as well as the ability of these clones to form both spontaneous (from the mammary fat pad) and experimental (tail vein injected) metastasis. For primary and spontaneous metastasis models, MOCK-2, M1 or M6 CA-IX MCF-7 cells were implanted in the mammary fat pads of mice, and growth was monitored by caliper measurement. Although *in vitro*, mock cells proliferated faster (Supplementary Fig. 7), this was not observed *in vivo,* as the tumors from CA-IX expressing clones grew significantly faster. At all of the time points measured, primary tumor volume was significantly increased in the CA-IX expressing clones, compared to controls (Fig. 4A **&** Supplementary Fig. 13). Regression-based analysis determined the growth rate of tumors was significantly different between the groups, p<0.0001. After resection, the primary tumors were sectioned, stained with H & E, and blindly graded by a board-certified pathologist (M.B.), who identified increased stromal invasion in the CA-IX expressing tumors (Fig. 4B). The mice were monitored for an additional 12 weeks post-resection, at which time the animals were sacrificed, and lungs stained for evidence of micro and macro metastases, scored blindly by a board-certified pathologist (M.B.). As shown in Table 1, 6/21 mice developed spontaneous metastasis from the M6 clone, whereas no spontaneous metastases were formed form the M1 clone and 1/10 were formed in the MOCK clones.

**Figure 4:**
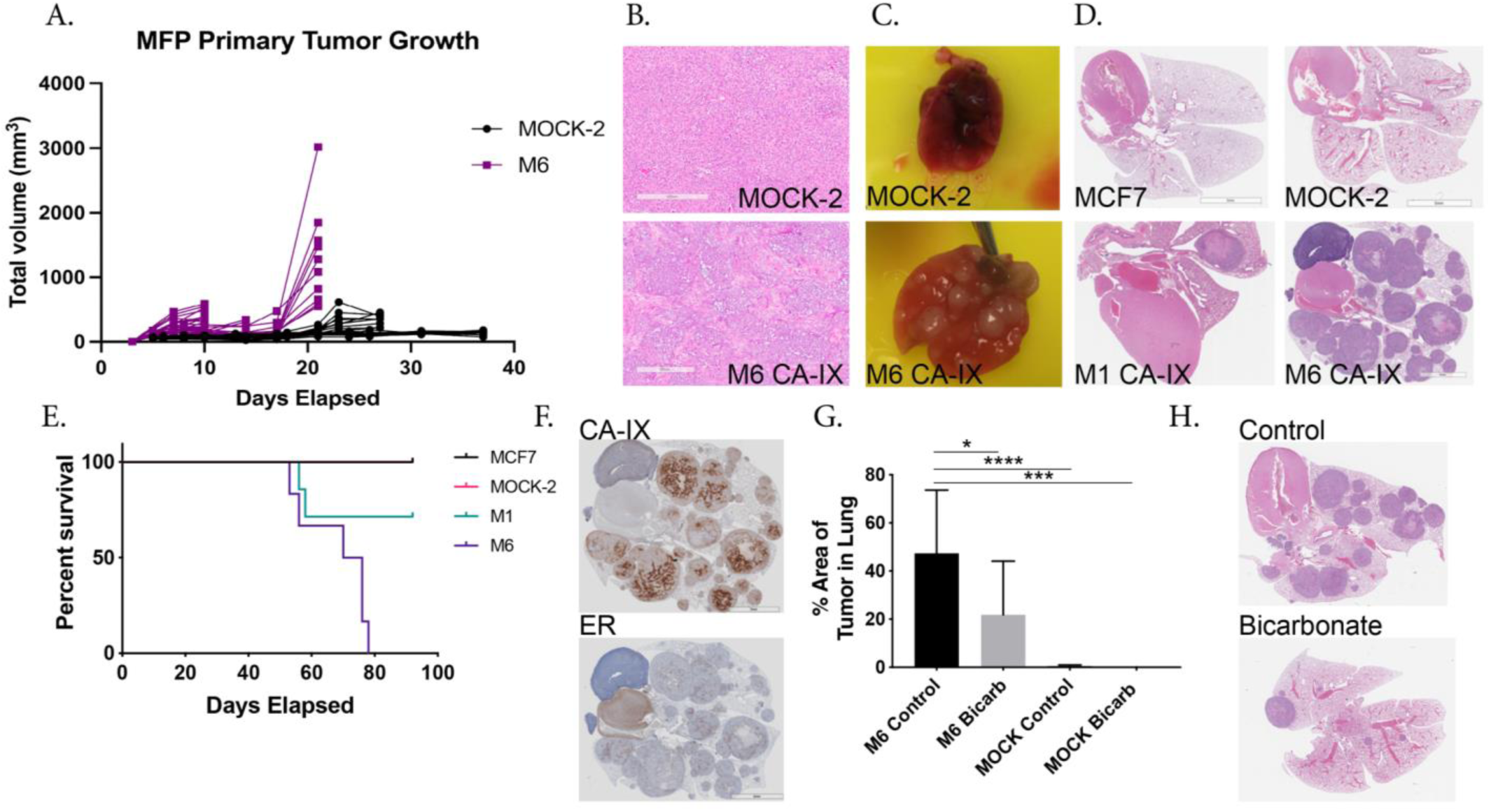
*In vivo* studies of CA-IX expressing cells and the effect on tumor growth, metastasis, and buffer therapy. **A:** Primary tumor volume of control or CA-IX cells implanted in the mammary fat pad, n=10 mice per group, statistical analysis using Welch’s T-test. Statistical analysis of the tumor volume was carried out to determine if tumor growth rate was different between the Mock-2 and M6 groups. Linear regression analysis showed the differences between the slopes were extremely significant. F = 21.03. DFn = 1, DFd = 116, P<0.0001 **B:** Representative H&E staining of resected primary tumors. M6 CA-IX group showed high infiltration of stromal cells compared to MOCK-2. **C:** Representative whole lung images from MOCK-2 and M6 experimental metastasis model showing the extent of metastasis visible. **D:** Representative immunohistochemistry of H&E staining in the lungs of the experimental metastasis groups, showing gross metastasis of the lungs. **E:** Kaplan-Meier Survival curve of experimental metastasis study in SCID beige mice, 90-day endpoint after tail vein injection of cells. Log-rank test performed p=0.0062, df=3. N=5(MCF-7), 6(MOCK-2), 7(M1), 7(M6) mice. **F:** Representative immunohistochemistry of M6 lungs with antibodies towards CA-IX and ER, to confirm the generated M6 clone were the cells forming tumors in the lungs. MCF-7 cells are ER-positive. **G:** Effect of buffer therapy on experimental metastasis of CA-IX clone M6 and MOCK-2 cells in SCID beige mice and the % of tumor burden in the lungs, 77days after IV injection of cells. Data are shown as average % ± SD, N=6(M6), 9(M6 Bicarb), 9(MOCK-2), 7(MOCK-2 Bicarb) mice, statistical analysis using ordinary one-way ANOVA. **H:** Immunohistochemistry of H&E staining in the lungs of the M6 control and bicarb treated experimental metastasis study groups, showing gross metastasis of the lungs. *p<0.05, **p<0.01, ***p<0.001, ****p<0.0001

**Table 1:**
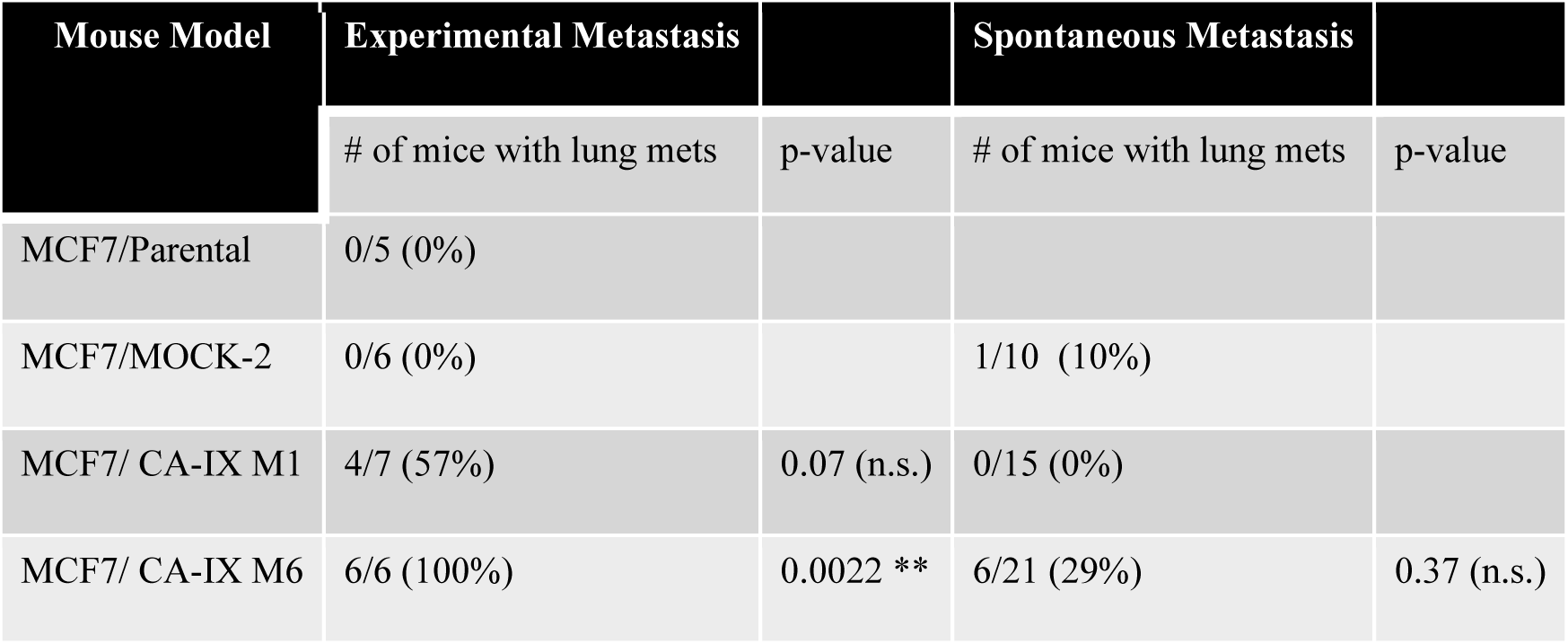
Effect of CA-IX expression on metastasis of MCF-7 cells. (Two-tailed Fisher’s exact t-test p<0.05*)

Because formation of spontaneous metastases is a multi-step, time-consuming, and complex process, we also investigated the ability of these clones to form experimental metastases following tail-vein injection, which only involves the final steps of extravasation and colonization. Metastases to the lung were scored blindly by a board-certified pathologist (M.B.), who observed that neither of the control groups developed metastases, and that both CA-IX clones had significant macrometastases (Table 1). Consistent with the spontaneous model, the M1 formed fewer metastases compared to the M6, however both CA-IX clones exhibited gross metastasis to the lungs (Fig. 4C), which were confirmed histologically (Fig. 4D). The resulting macrometastases significantly reduced overall survival of the mice (Fig. 4E). Additional IHC staining confirmed the lung metastasis expressed both CA-IX and human estrogen receptor, which we used as a marker for MCF7 cells (Fig. 4F).

Prior studies have shown that neutralization of acidity using oral buffers inhibits metastasis (Ibrahim-Hashim et al., 2017; Ibrahim-Hashim et al., 2012). As we hypothesized that our M6 CA-IX clones were metastatic by virtue of increased acid production (Fig. 1D, E), we asked whether buffer therapy would reduce the metastatic burden. Using the experimental metastasis model, we compared untreated to buffer-treated M6 or MOCK-2 mice. As in the first experimental metastasis study, all mice injected with M6 developed macrometastases, and these were significantly reduced by bicarbonate (Fig. 4G **&** H). Two out of ten MOCK-2 mice developed very small micrometastasis, and no metastases were found in the buffer therapy MOCK-2 group (Fig. 4G).

### Over-expression of yeast proton pump PMA1 increases motility and metabolism

While we hypothesize that CA-IX is acting as a proton equivalent exporting system, there are many other activities of this protein, including non-enzymatic activities, that could be activating glycolysis and promoting metastasis. To test whether the observed effects on glycolysis could be due to increased proton export, we utilized another model, PMA1, which electrogenically pumps H^+^ out of cells at the expense of ATP (Ferreira et al., 2001). Prior work has shown ectopic expression of PMA1 in murine 3T3 fibroblasts led to tumorigenesis (Perona and Serrano, 1988) and to increased aerobic glycolysis with elevated intracellular pH (Gillies, 1990; Martinez et al., 1994). We engineered MCF-7 cells to express PMA1 and selected two clones following zeocin selection (PMA1-C1 and PMA1-C5) as well as an empty vector transfected control (MOCK-1). PMA1 over-expression in C1 and C5 was confirmed, by qRT-PCR (Supplementary Fig. 14) and western blot (Fig. 5A). Moreover, using immunocytochemistry of non-permeabilized cells (Fig. 5B **&** Supplementary Fig. 15) or permeabilized cells (Supplementary Fig. 16), we verified PMA1 protein expression on the plasma membrane and in the cytoplasm of both PMA1 clones.

**Figure 5:**
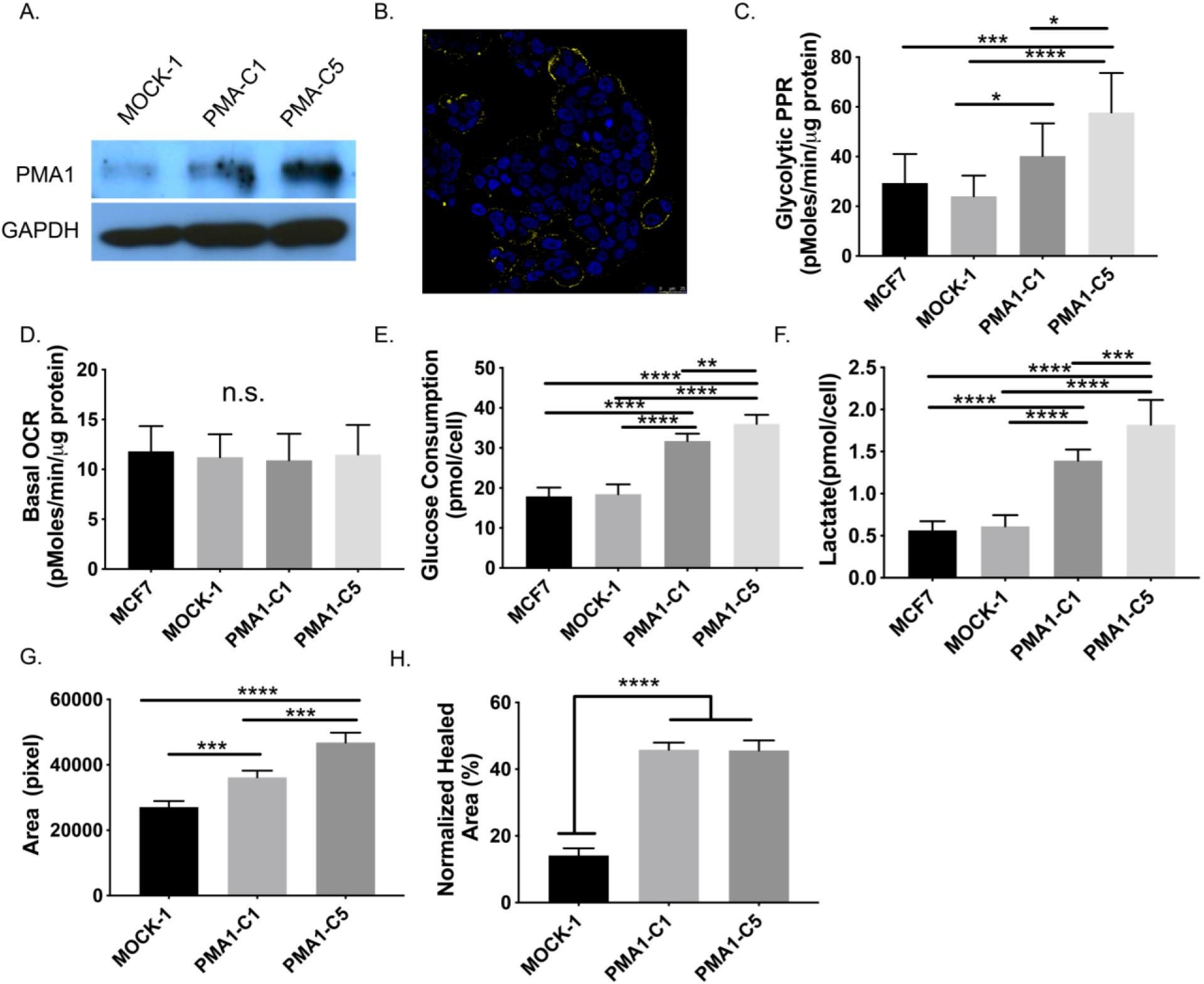
Over-expression of yeast ATPase proton pump, PMA1, in MCF-7 breast cancer cells increases glycolytic metabolism, migration, and invasion in vitro. A: Immunoblotting of protein lysates from MCF-7 cells transfected with empty vector (MOCK-1) or PMA1 vector (C1 and C5). Proteins from total cell extracts were immunoblotted for PMA1 and GAPDH (loading control). B: Representative immunocytochemistry image of PMA1 expression(yellow), overlayed with DAPI nuclear stain(blue), in PMA1-C5 non-permeabilized MCF-7 cells. PMA1 clones exhibit PMA1 staining, whereas MOCK-1 cells do not. C: Seahorse extracellular flux analysis of glycolysis associated proton production rate, measured after glucose is added, in MCF-7 parental, MOCK-1, PMA-C1, and PMA1-C5 MCF-7 cells. Data shown as mean ± SD n=9 biological replicates. Statistical analysis using ordinary one-way ANOVA. D: Seahorse extracellular flux analysis of basal oxygen consumption in the presence of glucose for MCF-7 parental, MOCK-1, PMA-C1, and PMA1-C5 MCF-7 cells. Data shown as mean ± SD n=20 biological replicates. Statistical analysis using ordinary one-way ANOVA. E: Glucose concentration in tumor-conditioned media collected after culturing to confluence from MCF-7 parental, MOCK-1, PMA-C1 and PMA1-C5 MCF-7 cells for 24hrs, measured using hexokinase activity assay. Data shown as mean ± SD n=9 biological replicates. Statistical analysis using ordinary one-way ANOVA. F: Lactate concentration in tumor-conditioned media collected after culturing to confluence from MCF-7 parental, MOCK-1, PMA-C1 and PMA1-C5 MCF-7 cells for 24hrs, measured using fluorescent lactate assay. Data shown as mean ± SD n=9 biological replicates. Statistical analysis using ordinary one-way ANOVA. G: Gel escape assay to measure migration and invasion in MOCK-1, PMA-C1 and PMA1-C5 MCF-7 cells. Cells are embedded in a Matrigel droplet without serum, and they are surrounded by media containing serum. Droplets are monitored over one week for cell invasion out of the droplet and the area measured is normalized to cellular proliferation rates. N=4 biological replicates. Statistical analysis using ordinary one-way ANOVA. H. Circular wound healing assay of MOCK-1, PMA-C1 and PMA1-C5 MCF-7 cells. A circular wound is created with a rubber stopper and then cells are monitored migrating into the area to close the wound. The area healed is quantified in % relative to starting area and normalized to the cellular proliferation rate. N=4 biological replicates. Statistical analysis using ordinary one-way ANOVA. *p<0.05, **p<0.01, ***p<0.001, ****p<0.0001

To characterize the metabolic activity of PMA1 transfectants, we again utilized the glycolysis and mitochondrial stress tests (GST and MST, respectively) of the Seahorse (XFe) Analyzer. The glucose-induced PPR (Fig.5C), and the glycolytic reserve (Supplementary Fig. 17) were significantly higher in the PMA1 clones compared to empty vector MOCK-1 or parental clones, suggesting functional activities of the transfected pump. In contrast to the CA-IX transfectants, there were no significant differences in oxygen consumption rates (OCR) between PMA1 clones and controls (Fig. 5D). This could be due to increased energy demand from the ATPase proton pump. We further confirmed these metabolic alterations by measuring glycolytic flux. PMA1 clones had significantly higher glucose consumption rates (Fig. 5E) and lactate production rates (Fig. 5F) compared to MOCK-1 or parental MCF-7 clones. These data, together with the CA-IX results, indicate that acid export can drive cells to exhibit a Warburg phenotype.

As with the CA-IX transfectants, we also measured invasion and migration using gel escape and circular wound healing assays, respectively. Compared to the MOCK-1 cells, both PMA1 clones expanded significantly more out of the gel drop (Fig. 5G **&** Supplementary Fig. 18). In the circular “wound-healing” assay, we monitored the migration of cells into a cell-free area. Again, compared to MOCK-1, both PMA1 clones had increased migration rates (Fig. 5H **&** Supplementary Fig. 19). Together, these results indicate that cellular invasion and migration were significantly enhanced by PMA1 expression and acid production.

### PMA1 induces a metastatic phenotype in vivo and alters the expression of proteins involved in metabolism and pH regulation

To investigate if proton export enhanced aggressiveness, as seen in the CA-IX model, we measured the PMA1 cells’ metastatic ability *in vivo* in both spontaneous and experimental metastasis models. In our *in vitro* studies, the proliferation rates of PMA1-C1 and the empty vector MOCK-1 clones were similar, whereas the growth rate of PMA-C5 was significantly slower (Supplementary Fig. 20). Hence, we omitted PMA-C5 in our *in vivo* studies to reduce the possibility of proliferation being a confounding variable. In both the spontaneous and tail vein metastases models, only 1 of 10 mice in each MOCK-1 group developed lung metastases. In contrast, 7 of 12 PMA-C1 mice developed lung metastases following tail vein injection and 4 of 9 formed spontaneous metastases (Supplementary Table 2.).

Lung metastases of PMA1 cells, visualized by H&E staining (Fig. 6A), were further validated by immunohistochemistry (IHC) of PMA1 (Fig. 6A) and RNA analysis of FFPE lung tissue for PMA1 gene expression (Supplementary Fig. 21). Notably, primary tumors revealed no significant growth differences between PMA1 and MOCK-1 tumors (Fig. 6B) or final tumor volume (Fig. 6C). However, blind grading (1 to 4+) of H&E-stained tumor sections by a board-certified pathologist (A.L.) indicated that PMA1 primary tumors were of significantly higher grade compared to MOCK-1 controls (p=0.016). The average grade was 2.7 ± 0.52 for MOCK-1 compared with 3.4 ±0.67 for the PMA1 primary tumors (Fig. 6D and 6E). Maintenance of PMA1 expression *in vivo* was confirmed with quantitative IHC of the resected primary tumors, demonstrating a significant difference in PMA1 protein expression between the MOCK-1 (19%± 3.0, n = 10) and PMA-C1 (90%± 2.5, n = 9) tumors (Fig. 6F **&** 6I).

**Figure 6:**
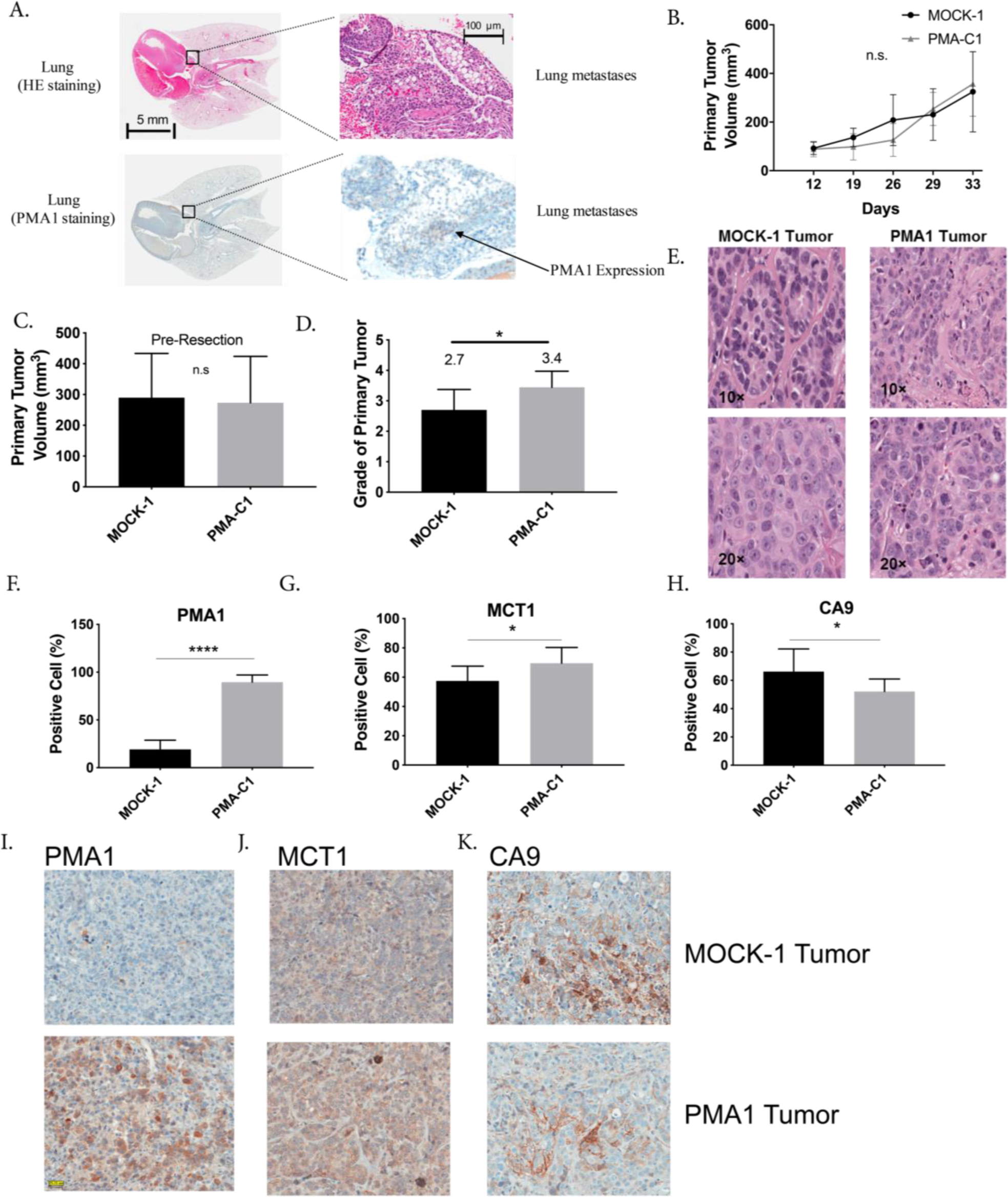
*In vivo* studies of PMA1 expressing cells and the effect on tumor growth, metastasis, and expression of metabolic markers. **A:** Representative lung images (5mm and 100um) from the experimental metastasis study in SCID beige mice, whereby MOCK-1 or PMA-C1 cells were injected via the tail vein and allowed to grow for 3 months. At the endpoint lungs were resected and sections were stained with H & E to look for metastases and PMA1 to confirm expression in the PMA C1 metastases. **B:** Primary tumor growth rate in SCID beige mice of MOCK-1 or PMA1-C1 cell lines, injected subcutaneously. Data are shown as mean ± SD over time, N=7(MOCK-1) N=6 (PMA1-C1). **C:** Quantification of primary tumor volume, MOCK-1 or PMA1-C1, resected after 33 days of growth in SCID beige mice. Tumors were resected to allow for spontaneous metastasis studies to continue. Data are shown as mean ± SD, N=9 (MOCK-1) and N=9 (PMA1), statistical analysis using Unpaired t-test with Welch’s correction. **D:** Histological grade of MOCK-1 and PMA1 tumors. Data are shown as mean ± SD, N=10 (MOCK-1) and N=9 (PMA1), statistical analysis using Unpaired t-test with Welch’s correction. **E:** Representative images of H & E staining in MOCK-1 and PMA1 primary tumors were used to score histological grade. **F.** Quantification of immunohistochemistry staining for PMA1 protein in FFPE sections of resected primary tumors, MOCK-1 and PMA1-C1. Data are shown as mean ± SD, N=10 (MOCK-1) and N=9 (PMA1), statistical analysis using Unpaired t-test. **G:** Quantification of immunohistochemistry staining for MCT1 protein in FFPE sections of resected primary tumors, MOCK-1 and PMA1-C1. Data are shown as mean ± SD, N=10 (MOCK-1) and N=9 (PMA1), statistical analysis using Unpaired t-test with Welch’s correction. **H:** Quantification of immunohistochemistry staining for CA-IX protein in FFPE sections of resected primary tumors, MOCK-1 and PMA1-C1. Data are shown as mean ± SD, N=10 (MOCK-1) and N=9 (PMA1), statistical analysis using Unpaired t-test. **F:** Immunohistochemistry representative images of PMA1 and MOCK-1 primary tumors stained for PMA1 protein expression. **I:** Representative immunohistochemistry images of FFPE sections of resected primary tumors, MOCK-1 and PMA1-C1 stained with antibody for MCT1. **K:** Representative immunohistochemistry images of FFPE sections of resected primary tumors, MOCK-1 and PMA1-C1, stained with antibody for CA-IX. *p<0.05, **p<0.01, ***p<0.001, ****p<0.0001

Additional IHC of PMA1 tumors showed that they had significantly lower CA-IX expression than MOCK-1 (Fig. 6H **&**6K). As CA-IX plays a vital role in regulating tumor pH (Doherty et al., 2014; Lee et al., 2018; Swietach et al., 2009), we postulate that its activity may have been made redundant by PMA1. PMA1 expressing tumors also had higher levels of the monocarboxylate (lactate) transporter, MCT1 (Fig. 6G **&**6J), which has been associated with increased aggressiveness in breast cancer (Doherty et al., 2014). Notably, other proteins, such as glucose transporter 1, GLUT1 (Supplementary Fig. 22), the sodium hydrogen exchanger 1, NHE1 (Supplementary Fig. 23), and MCT4 (Supplementary Fig. 24) showed no differences between the PMA1 and MOCK-1 groups. These data suggest that the increased glycolytic flux, which requires glucose uptake by GLUT1, can be accommodated by native protein levels of GLUT1 (i.e., it is not rate-limiting).

## DISCUSSION

The primary goal of this study was to determine if acid export *per se* could drive aerobic glycolysis, the “Warburg Effect”, in cancer. Aerobic glycolysis, a cancer hallmark, is often associated with more aggressive tumors. Numerous studies have attempted to determine why tumors favor fermentative glycolysis, even in the presence of sufficient oxygen (Gatenby and Gillies, 2004a; Vander Heiden et al., 2009). Aerobic glycolysis is bioenergetically inefficient, producing only 2 ATP per glucose, compared to 36 ATP upon complete oxidation. Thus, it is not obvious why this should be such a prevalent phenotype. It is axiomatic that common phenotypes *must* confer a selective advantage, yet the evolutionary drivers of aerobic glycolysis are not clear. Theories proposed include glucose addiction (Buzzai et al., 2005), dysfunctional mitochondria (Shiratori et al., 2019), which is not the case in most cancers, and to meet the rapid energetic requirements of membrane transporters (Epstein et al., 2014). Another theory is that glycolysis enhances tumor cell proliferation by generating the anabolic building blocks for macromolecules. However, most of the glucose-derived carbons are exported out of the cell as lactate (Lunt and Vander Heiden, 2011). Moreover, TCA cycle intermediates are considered more important for lipid, amino acid, and nucleotide synthesis, compared to glycolytic intermediates (Carracedo et al., 2013; Wellen and Thompson, 2012). However, none of these theories clearly demonstrate why so many cancers favor aerobic glycolysis. We hypothesize that acid export *per se* and extracellular acidification provides a distinct selective advantage, and that it is enabled by increased glucose fermentation. This theory was first proposed as the “acid mediated invasion hypothesis” (Gatenby et al., 2006a) and has been subsequently elaborated (Gillies et al., 2008).

To test the hypothesis that acid export increases glycolysis in cancer, we over-expressed a proton exporter, CA-IX, in an OXPHOS dominant cell line, MCF-7. In MCF-7 cells, CA-IX is not expressed under normoxic conditions, and over-expression resulted in cells that more rapidly exported acid and up-regulated glycolysis. Both glucose consumption and lactate production rates increased. Other studies have shown that CA-IX over-expression can increase lactate production, albeit under hypoxic conditions (Jamali et al., 2015). As glycolysis was specifically increased, we investigated the mechanisms whereby acid export could be causing this shift in metabolism. We broadly interrogated cellular metabolism using a targeted metabolomics panel and found that the most significantly altered metabolites in the CA-IX expressing clones were glycolytic intermediates. Specifically, all glycolytic intermediates upstream of pyruvate kinase were increased. Enzyme activity can be affected by pH, including those in the glycolytic pathway, and are most active at the pH of their subcellular compartment from acidic lysosomes to alkaline mitochondria (Persi et al., 2018). For glycolytic enzymes, pH optima are slightly on the alkaline side of neutral (7.2-7.4), meaning that raising pH above neutrality will globally increase activities of glycolytic enzymes. Because the data suggested pleiotropic increases in enzyme activity, we measured intracellular pH using fluorescence ratio imaging in our CA-IX expressing clones. These clones, had a higher pHi in biologically relevant extracellular pH conditions, compared to the parental. Specifically, at pHe 6.8 and 7.2, at which CA-IX can function as a proton equivalent exporter, the CA-IX clones had increased pHi compared to parental. One caveat of this experiment was that the MOCK cells were unquantifiable as they did not accumulate SNARF-1, possibly due to increased activity of multidrug resistance transporters. In addition, the experiments in low-chloride directly implicate increased pHi in regulating aerobic glycolytic flux. These data indicate loading cells with HCO_3_^-^ ions raises the intracellular pH sufficiently to enhance enzyme activity and result in increased glycolytic flux.

We tested our hypothesis using two more models and another acid exporting protein to minimize cell line and protein-specific effects. Similar to the MCF-7 results, over-expression of CA-IX in U2-OS and HEK293 cells increased glycolysis compared to controls. Although parental HEK293 cells are more glycolytic than the other cell lines chosen, over-expression of CA-IX still enhanced glycolysis. We tested another acid exporting protein, PMA-1, which has an unequivocal activity of exporting protons at the expense of ATP. PMA-1 over-expression in MCF-7 cells similarly resulted in increased glycolysis, as measured by increased glucose uptake and lactate production. These findings, together with our CA-IX results, suggest that expression of proton exporting activity can up-regulate aerobic glycolysis, likely through a global increase of intracellular pH. Notably, while the CA-IX transfectants had reduced oxygen consumption, this was not observed in the PMA1 cells. The cause of this difference is not known and may reflect differences in the bioenergetic requirements for the two transporting systems.

We hypothesize that acid export-driven glycolysis would make them more aggressive, as measured in vitro with motility and invasion assays, and in vivo with experimental and spontaneous metastases studies. Glycolysis and acidity have been correlated with poor prognosis and metastasis (Walenta et al., 2000; Webb et al., 2011a). Our focus on CA-IX as an acid exporter was due to its clinical relevance in many cancer types, such as breast, ovarian, and astrocytoma, where CA-IX over-expression correlates with poor prognosis, reduced survival, and reduced metastasis-free survival. This suggests CA-IX specifically and perhaps acid export generally enhances cancer aggressiveness and subsequently metastasis. In our models, *in vitro* acid export driven by CA-IX or PMA1 was linked to enhanced migration and invasion *in vitro*, which is consistent with prior studies (Csaderova et al., 2013; Shin et al., 2011; Svastova et al., 2012). *In vivo*, cells at the invasive edge tumor periphery are known to be more acidic and express CA-IX (Lloyd et al., 2016; Rohani et al., 2019). CA-IX has also been shown to enhance matrix metalloproteinase activity, in particular MMP-14 which is active at an acidic pH, resulting in stromal degradation that aids cancer migration into the periphery (Swayampakula et al., 2017).

Our encouraging *in vitro* results led us to take these models, CA-IX and PMA-1, *in vivo*. We found that aggression and metastasis were higher in both PMA-1 and CA-IX transfectants. Primary tumor growth was enhanced in the CA-IX model compared to controls. In the PMA-1 model, a pathologist blindly scored the primary tumors a higher grade compared to control tumors. However, spontaneous metastasis after primary tumor resection was not significantly increased in PMA-1 or CA-IX transfectants compared to controls (Table 1 **&** Supplementary Table 2). It is notable, however, that in our spontaneous models 10/45 mice with PMA-1 or CA-IX clones had metastases, compared to 2/20 control mice with parental or MOCK-transfections. In contrast, our experimental metastasis model, which skips the intravasation step, showed enhanced experimental metastasis in both models, with 17/25 mice in PMA-1 or CA-IX transfectants developing metastasis, compared to 1/21 metastases in control mice (Table 1 **&** Supplementary Table 2). We did not quantify the number or size of metastatic lesions, because the important metric is binary: i.e. whether or not these clones were able to metastasize at all. A related study in 4T1 breast cancer showed that inhibiting CA-IX reduced tumor growth and experimental metastasis (Lou et al., 2011). However, inhibition is different than induction, and 4T1 are highly glycolytic and acidic to begin with. However, this study does indicate the importance of acid export and its role in enhancing tumor growth. Due to the robust enhanced metastasis formation in the experimental metastasis studies, it indicates that acid export can facilitate tumor cell extravasation out of the blood vessels and colonization of metastatic sites.

In our CA-IX model buffer therapy significantly reduced tumor burden in the lungs compared to their untreated counterparts. Although this did not completely prevent metastasis, combinations of buffer therapy with specific acid exporter inhibitors may be necessary. CA-IX is minimally expressed in normal tissue and could be a viable therapeutic target (Silvia Pastorekova et al., 1997) and other proton pump inhibitors are currently in clinical trials (Fais, 2015). Many studies have hinted at the importance of acidity, and many have proposed reasons as to why cancer cells favor aerobic glycolysis, but few have proposed that acidity is the driver. This study represents the first to test whether acid export can increase aerobic glycolysis and enhance cancer aggressiveness, rather than acid merely being a by-product.

## Materials and Methods

### Construction of stable cell lines

#### Plasmids

Yeast plasma membrane ATPase 1 (PMA1) cDNA construct was designed based on the sequence (Accession Number: NM_001180873; Saccharomyces cerevisiae S288c PMA1). The codons were optimized for the suitable expression in mammalian cells and restriction enzyme sequences Hind III and Xho I were inserted at the 5’ and 3’ ends of the full-length sequence, respectively. The fully designed DNA sequence was commercially synthesized (Blue Heron Biotechnology, Bothell, WA, 98021). This was cloned into pcDNA3.1/Zeo (+) vector in which PMA1 gene expression was driven under the CMV promoter. The sequence of the pcDNA/PMA1 construct and the identity of the parental cell line were confirmed by the molecular genomics core facility (Moffitt Research Institute, Tampa, FL). Carbonic anhydrase 9 (CA9) construct was designed by Origene based on the sequence(Accession Number:NM_001216 ; Homo sapiens) and cloned into a pCMV6 vector (PS10001, Origene, MD) to form pCMV6/CA-IX vectors (CQ10630, RC204839 subclone, Origene, MD)in which CA9 gene expression was driven under the CMV promoter.

#### Cell lines

The MCF-7 cells and HEK 293 cells as transfection host cell lines were acquired from American Type Culture Collection (ATCC HTB-22, Manassas, VA) and maintained in RPMI media 1640 (Life Technologies Gibco®, 11875-093) supplemented with 10% FBS (Hyclone Laboratories, UT) under standard cell culture conditions. The U2-OS cell line was a gift from Jillaina Menth, Moffitt Cancer Centre Translational Research Core, and maintained in RPMI media 1640 (Life Technologies Gibco®, 11875-093) supplemented with 10% FBS (Hyclone Laboratories, UT) under standard cell culture conditions. The MCF-7 cells were transfected with empty pcDNA, pcDNA/PMA1, pCMV6 (PS10001, Origene, MD) and pCMV6/CA-IX vectors (CQ10630, RC204839 subclone, Origene, MD) respectively, resulted in MCF-7/MOCK-1 cells, MCF-7/PMA1, MCF-7/MOCK-2, and MCF-7/CA-IX cell lines by standard clonogenic stable cell construction procedures using Fugene HD (Promega, E 2311). The U2-OS and HEK-293 cells transfected with empty pCMV6 (PS10001, Origene, MD) and pCMV6/CA-IX vectors (CQ10630, RC204839 subclone, Origene, MD) respectively, resulted in U2-OS/MOCK-2, HEK/MOCK-2, and U2-OS/CA-IX and HEK/CA-IX clones. Briefly, a number of individual single clones were selected in the media containing 300 *µ*g/ml zeocin (Invitrogen, 450430, Carlsbad, CA), or 300 *µ*g/ml G418 and stable expression in individual clones were verified using western blotting. Cell lines were tested for mycoplasma using MycoAlert assay (Lonza).

### Spheroid formation

Spheroids were formed as previously described (Russell et al., 2017). Briefly, cells were suspended in Perfecta 3D hanging drop plates (HDP1096, 3D Biomatrix, MI) at 25K cells/40ul droplet. Spheroids were allowed to form for 5 days and then centrifuged at 450rpm, with no brake, into media-containing Costar Ultra low attachment U-bottom 96-well plates (CLS3474, Corning, NY). Spheroids were imaged in the Celigo Imaging Cytometer (Nexcelom Bioscience, MA) using bright field imaging single colony verification analysis.

### Western blotting

#### Chemiluminescence

The cell membrane protein samples were collected using Mem-PER eukaryotic membrane protein extraction reagent kit (Thermo Scientific, 89826, MA) according to the protocol instruction, and the protein samples were further purified and concentrated by Pierce SDS-PAGE sample prep kit (Thermo Scientific, 89888). Thirty micrograms of protein per sample was separated on polyacrylamide-SDS gels and electrophoretically transferred to nitrocellulose membranes. Membranes were incubated with primary antibody against PMA1 (1:1000, Abcam, ab4645), and GAPDH (1:1000, Santa Cruz Biotechnology, TX, se-25778). For visualization, horseradish peroxidase (HRP)-conjugated secondary antibodies: Goat anti-rabbit IgG HRP and goat anti-mouse IgG HRP, followed by ECL kit (Thermo Scientific, 32209) were used.

#### Fluorescence

Fifteen *µ*g of protein per sample was separated on a BioRad Mini-protein 4-15% precast 12 well 20 *µ*l gels (4561085, Bio-Rad, CA) and electrophoretically transferred to Odyssey Nitrocellulose membrane (P/N 926-31092, LI-COR, NE). Membranes were blocked with Odyssey TBS Blocking Buffer ( P/N 927-50000, LI-COR, NE) and incubated with primary antibody against CA-IX(1:1000, M75 mouse monoclonal CA-IX, Bioscience Slovakia), ER alpha ( 1:2000, Rabbit polyclonal ab 75635, Abcam), rabbit Anti-CA12 antibody [EPR14861] - C-terminal (ab195233), rabbit Anti-Carbonic Anhydrase II antibody [EPR5195] (ab124687), β-Actin(1:2000, (8H10D10) Mouse mAb #3700-Cell Signaling), GAPDH (1:4000, rabbit monoclonal ab 181602, Abcam). For visualization, IRDye Fluorescent secondary antibodies (LI-COR) were used: IRDye 680RD Goat Anti-mouse IgG (H+L), IRDye 680RD Donkey anti-rabbit IgG (H+L), IRDye 800CW goat anti-mouse IgG(H+L) and IRDye 800CW donkey anti-rabbit IgG(H+L). Membranes were imaged on LI-COR Odyssey Blot Imager and quantified using Image Studio Version 2.1(LI-COR). Uncropped versions of western blots are available in Supplementary.

### Immunocytochemistry

Cells were grown on glass coverslips and fixed in 4% paraformaldehyde (Sigma-Aldrich) for 10 min at room temperature. Cells were blocked in 5% BSA for 1hr at room temperature. Cells were stained with the PMA1 antibody (1:100; SC-33735, Santa Cruz Biotechnology) or CA-IX antibody for 2 hours (1:500, ab184630, Abcam) and washed in PBS. Cells were further incubated for 1 h with secondary anti-rabbit-Alexa Fluor 594 antibody (1:2000; A11072, Invitrogen) or anti-mouse-Alexa Fluor 594(1:2000; A11005, Invitrogen) and additionally incubated in WGA, cell membrane marker (W6748, Invitrogen) for 10 min on ice. The cells were mounted for fluorescence with DAPI (H-1200, Vector). The slides were viewed by Leica inverted SP5 AOBS confocal microscope, and micrographs were taken, and images were subsequently acquired in the Moffitt Analytic Microscopy Core Facility by using dual photomultiplier tube detectors and LAS AF software (Leica Microsystems). For detection of intracellular PMA1, cells were fixed and permeabilized with 1:1 mixture of methanol and acetone, and immunostained with PMA1 antibody for 1h, followed by 1 hour of incubation with the secondary anti-rabbit Alexa488 antibody (Molecular Probes, Invitrogen). The cells were mounted and viewed by fluorescence microscopy.

### Proliferation rate assay

Cells were cultured in a 24-well plate under standard cell culture conditions for 24, 48, 72, 96 hours, and the cell number and viability were determined with a trypan blue dye by using the Countess automated cell counter (Invitrogen).

### Oxygen consumption and proton production rate measurements (OCR and PPR)

Real-time oxygen consumption (OCR) and proton production rate (PPR) were measured by using the Seahorse Extracellular Flux (XFe-96) Analyzer (Seahorse Bioscience, Chicopee, MA). The cells were seeded in an XFe-96 microplate (Seahorse, V3-PET, 101104-004) in normal growth media overnight. The growth media were replaced with DMEM powder base media ( Sigma D5030) supplemented with 1.85g/L sodium chloride and 1mM glutamine, and the cells were incubated in the media in the absence of glucose, when testing glycolysis, in a non-CO_2_ incubator for one hour prior to the measurement. PPR and OCR were measured in the absence of glucose associated with the non-glycolytic activity, followed by two sequential injections of D-glucose (6mM) and oligomycin (1*µ*M) in real-time, which are associated with glycolytic activity and glycolytic capacity (reserve). The mitochondrial stress test was also used where cells were incubated in glucose (5.5mM), and glutamine (1mM) containing media and basal OCR and PPR measured, prior to sequential injection of Oligomycin (1 *µ*M), associated with ATP linked OCR, FCCP(1*µ*M) associated with mitochondrial reserve capacity and Rotenone/Antimycin A (1*µ*M). Following the measurements, protein concentrations were determined *in situ* for each well using a standard BCA protein assay (Thermo Scientific Pierce). The OCR and PPR values were normalized to *µ*g protein. Results were also normalized using Celigo High Throughput Micro-Well Imaging Cytometer (Nexcelom Bioscience) by bright-field direct cell counting and normalized per 10K cells prior to assay.

### Glucose consumption and lactate production assays

Figure 1: Cells were seeded in a 96-well plate in the growth media containing 10% FBS. Once cells reached 90% confluence, the growth media were removed, and the cells were washed twice in PBS and media was replaced for 24 h. The media were collected from 24 h incubation for both glucose consumption and lactate production assays. The cells were then trypsinized and the cell densities were determined. Glucose quantification was conducted using glucose bioluminescent assay kit (Glucose Glo Assay, Promega) as described per manufacturer instruction. The lactate assay kit (Lactate Assay kit, Sigma Aldrich) was used to measure L (+)-Lactate in the culture media according to the manufacturer’s instructions. Data were normalized by cell density per well and were reported as lactate production and glucose consumption per ug protein.

Figure 5: Cells were seeded in a 6-well plate in growth media containing 10% FBS. Once cells reached 90% confluence, the growth media were removed, and the cells were washed twice in PBS and incubated in serum-free and phenol-red free media for 24 h. The media were collected from 24 h incubation for both glucose consumption and lactate production assays. The cells were trypsinized and the cell densities were determined. Glucose quantification was conducted using glucose colorimetric/fluorometric assay kit (BioVision, K606-100) as described per manufacturer instruction. The lactate assay kit II (BioVision, K627-100) was used to measure L (+)-Lactate in the culture media according to the manufacturer’s instructions. Data were normalized by cell density per well and were reported as lactate production and glucose consumption as pmol per cell.

### Glucose uptake radioactive assay

Cells were seeded in 24 well plates to 80% confluence. Cells were incubated for 1hr with 1 *µ*Ci of Deoxy-D-glucose, 2-[1,2-^3^H(N)] (NET549250UC, Perkin Elmer, MA) at 37*◦*. Cells were washed 2 × DPBS and lysed with 300ul of NaOH, cell extract was added to a vial with 6ml of Eoscint XR scintillation liquid (LS-272, National Diagnostics, GA). Uptake was quantified by measuring ^3^H in a Perkin Elmer TriCarb scintillation counter and normalizing to protein concentration.

### YSI 2950D biochemical analysis of lactate

Cells were seeded in a 96-well plate in the growth media containing 10% FBS. Once cells reached 90% confluence, the growth media were removed, and the cells were washed twice in PBS and media was replaced for 24 h or 48hr. The media were collected from 24 h & 48hr incubation for lactate production measurement. The cell densities per well were determined by Celligo imaging cytometer brightfield cell count application. Lactate quantification was measured by the YSI. Data were normalized by relative cell number and were reported as lactate production in g/L/per cell.

### Untargeted metabolomics

Samples were prepared according to Beth Israel Deaconess Medical Centre Mass Spectrometry Core and run on a Thermo QExactive Plus/HF Orbitrap LC-MS/MS. Briefly, cells were grown to 80% confluence in 10cm^2^ dishes. Cells were changed into fresh media two hours prior to collection, media was aspirated, cells were washed in ice cold PBS, and 1ml of 80% methanol (−80°C) added to plate on dry ice, then transferred to -80°C freezer for 15 minutes. The cell plate was scraped on dry ice and contents collected. Sample was spun in cold centrifuge at max speed for 20min and supernatant removed. Supernatant was dried in speed vac for 5hrs then stored at -80°C. Before Mass Spec analysis, samples were resuspended in HPLC grade water relative to protein concentration. Data were analyzed using Metaboanalyst online software, there was no data filtering, and data were normalized by sum of all metabolites per sample.

### Intracellular pH (pHi)

#### Solutions and media

(i) Solutions for Seahorse experiments: 2mM HEPES, 2mM MES, 5.3 mM KCl, 5.6 mM NaPhosphate, 11 mM glucose, 133 mM NaCl, 0.4 mM MgCl_2_, 0.42 mM CaCl_2_, titrated to given pH with NaOH. For reduced Cl^-^ experiments, 133 mM NaCl was replaced with 133 mM NaGluconate and MgCl_2_ and CaCl_2_ were raised to 0.74 mM and 1.46 mM, respectively, to account for gluconate-divalent binding. Calibration solutions for nigericin: 145 mM KCl, 1 mM MgCl2, 0.5 mM EGTA, 10 mM HEPES, 10 mM MES and adjusted with NaOH to required pH. pHe media to measure pHi: Solution A: 125mM NaCl, 4.5mM KCl, 1mM CaCl_2_, 1mM MgCl_2_, 11mM glucose base media. Split solution A into two parts-Solution B: 22mM HCO_3_^-^ and Solution C: 22mM NaCl. Mix B&C as follows-40ml B & 0ml C= 22mM HCO_3_^-^ (pH7.41), 20ml B & 20ml C =11mM HCO_3_^-^ (pH7.11), 10ml B & 30ml C=5.5mM HCO_3_^-^ (pH6.81), and 4ml B & 36ml C=2.2mM(pH6.41).

#### Fluorescent Labeling and Image Analysis

Cells were loaded with 10uM cSNARF1 and 2.7uM Hoechst 33342 for 10mins at 37°. Cells were washed in neutral pH media and imaged in various pHe media (6.4-7.4) on Leica SP5 Confocal Microscopy x40 objective. Cytoplasmic pH was measured by gating pixels according to a threshold level of Hoechst signal within cSNARF1-positive pixels. Fluorescence at 580 and 640 nm was averaged, background offset and ratioed for each particle representing a cell.

### Cell invasion and migration assay in vitro

*In vitro* cell motility and invasiveness were measured by methods as previously reported with some modifications _30_. The motility change was measured by the circular wound healing assay using OrisTM Cell Migration Assay Kit (Platypus, CMAU101). Cells were plated on a 96-well plate at 1 × 10^6^ cells/ml while a cell seeding stopper masker the circular area at the center of each well. The cell seeding stoppers were removed 24 hours after the plating and cells were cultured a further 30 hours to monitor the closing of the cell-free area (wound area). The area covered by live cells was measured by labeling cells with Calcein-AM (Life Technology, C3099) and analyzing microscopic images (2.5x) by Image J (NIH). The smaller wound size represents the higher motility.

Wound healing scratch assay was used to measure motility into CA-IX clones. 96-well plates were seeded with 1 × 10_6_ cells/ml per well in 10% FBS, 1% PenStrep RPMI-1640 and incubated overnight. The plate was uniformly scratched with Essen Bioscience 96 Woundmaker, media was removed, and the plate washed with DPBS and then 200ul of media was added to each well. The wound was imaged on Celigo Imaging Cytometer and the number of cells that migrated into the wound was quantified by direct bright field cell counting at 0, 24, 48hr.

Cell invasiveness was measured by monitoring cells escaping from Matrigel (Becton Dickenson, 356231). Cells were suspended in 50% Matrigel in serum-free RPMI-1640 at 1 × 10_7_ cells/ml. A Matrigel droplet (volume = 5 *µ*l) was placed on each well of a 24-well plate. The Matrigel was solidified by incubating at 37*^°^*C overnight and then FBS-containing normal growth media was added. The cells escaped from the Matrigel droplet were monitored in real-time by using the IncuCyte ZOOM system (Essen BioScience) or Celigo Imaging Cytometer (Nexcelom Bioscience, MA). After 7 days of culture, the cell expansion from the droplets was quantified by Celigo single colony verification algorithm or Image J after fixing cells in 3.7 % formaldehyde and staining in crystal violet solution. The larger area occupied by cells represents the higher invasion potential.

#### Normalization of invasion and migration assays

The results of invasion and migration assays were normalized by the proliferation rates of the cells. Proliferation rates were calculated by a linear fit of cell growth, 48 hours after seeding the cells (See Figure S16) and weighted by the standards error. The normalization was carried out in order to eliminate inherent differences between clones making it possible to compare them. In the case of the invasion assay, final growth was divided by the proliferation rate of each of the cells. For the migration assay (i.e. wound healing assay), the relative healed area was divided by its corresponding growth rate and then multiplied by the growth rate of the MOCK-1 cells. This allowed comparison between the normalized healed areas of each clone, to one of the MOCK-1 cells. We used the Matlab R2012a curve fitting toolbox (The MathWorks, MA). Matlab code is available in the Supplementary materials.

### SCID mice

Six-week-old female SCID Beige mice were purchased from Charles River Laboratories. Mice were given a week to acclimate to the animal facility before they were studied. To minimize the risk of any exogenous infection, the SCID mice were maintained and cared for in a sterile, static micro-isolation cage. Mice received irradiated food (Harlan Laboratories) and sterile water *ad libitum*. All animal experiments were performed under a protocol approved by the University of South Florida Institutional Animal Care and Use Committee.

### Metastasis assays in vivo

Since MCF-7 cells are estrogen-dependent for tumor formation, estrogen pellets, 17 *β* estradiol, 0.72 mg/pellet, 60-day release (Innovative Research of America, SE-121) were subcutaneously implanted in the shoulder region of the mice two days prior to tumor inoculation. For the primary tumor growth study with MCF-7/MOCK-2 or MCF-7/ CA-IX M6 cells, mice were given 200nM 17 *β* estradiol in drinking water to try and prevent adverse side effects of pellet use, including balder stones and urinary tract infections.

For the experimental metastasis, SCID mice were injected through tail veins with 1 *×* 10^6^ cells in 200 *µ*l of PBS solution (either MCF-7/MOCK-1, MCF-7/PMA1-C1, MCF-7/MOCK-2, MCF-7/CA-IX M1 or MCF-7/CA-IX M6 cells). Three months after injection of the cancer cells, the mice were euthanized, and the lung tissues were surgically excised, fixed, and stained with hematoxylin and eosin (HE). Lung sections (at least three histologic sections for each lung specimen) were examined for metastatic nodules under a light microscope by a breast cancer pathologist (A.L or M.M.B) who was blinded to identifiers.

For the spontaneous metastasis studies, approximately 10*×*10^6^ cells (either MCF-7/MOCK-1 and MCF-7/PMA1-C1) in 100 *µ*l of PBS +100 *µ*l of Matrigel were injected into the mammary fat pads of mice. Once tumors reached approximately 400 mm^3^, or 6 weeks post-cell injection, the tumors were resected, fixed, and stained with H&E, or PMA1 antibody. Three months after resection, the mice were sacrificed, and lung sections were examined for lung metastases. The tumors were measured twice every week throughout the study with a digital caliper and volume values were calculated with the formula V= (Length*×*Width^2^)/2. The body weights were monitored twice a week throughout the study.

### Treatment model

Female SCID Beige mice received 200mmol/L of sodium bicarbonate water 2 days prior to tail vein injection for experimental metastasis study. Mice continued receiving bicarbonate water until the end of the experiment. Control mice received regular tap water. Since MCF-7 cells are estrogen-dependent for tumor formation, estrogen pellets, 17 *β*-estradiol, 0.36 mg/pellet, 90-day release (Innovative Research of America, SE-121) were subcutaneously implanted in the shoulder region of the mice two days prior to tumor inoculation. For the experimental metastasis, SCID mice were injected through tail veins with 1 *×* 10^6^ cells in 200 *µ*l of PBS solution (MCF-7/MOCK-2 and MCF-7/CA-IX M6 cells). Once MCF-7/CA-IX M6 control group had observable lung metastasis by T2 MRI (∼74 days), mice were humanely euthanized, and lungs and kidney collected.

### Histology

The tissues were harvested, fixed in 10% neutral buffered formalin (Thermo Scientific), processed, and embedded in paraffin. Tissue sections (4 *µ*m) were prepared and stained with H&E in the Moffitt Cancer Center Tissue Core. The histological slides of resected primary breast tumor xenografts from MCF-7/MOCK-1 or MCF-7/PMA1-C1 groups, and MCF-7/MOCK-2, MCF-7/CA-IX M1 and MCF-7/CA-IX M6 groups. were blindly examined under a light microscope by a pathologist (A.L or M.M.B) for tumor grades using the most common grading system: G1: well differentiated (low grade); G2: moderately differentiated (intermediate grade) G3: poorly differentiated (high grade); G4: undifferentiated (very high grade), as assessed according to histological features of stromal hypercellularity, atypia, stromal mitotic activity, presence of stromal over-growth and mitosis, necrosis, spindle cells differential and chromatin activity. Tumor burden in the lung was measured by Aperio ImageScope (Leica Biosystems, IL) and calculated as % of total lung tissue and compared between groups. Tumors were drawn around by hand using the Aperio software and confirmed by a pathologist (M.M.B), % area of tumor in lungs was then calculated by comparing the area of total lung tissue to the area of the tumor within lungs. For all lung metastases IHC, three different sections were taken from each lung and analyzed, with 5-6 sections taken between each analyzed slice to ensure entirety of lungs, and metastatic burden was analyzed.

### Immunohistochemical (IHC) staining

The cross-sections were stained with various antibodies as per normal laboratory protocol in the Moffitt Tissue Core Histology Facility. Positive and negative controls were used for each antibody staining and staining condition was optimized for each antibody. The antibodies were utilized in this study as follows: rabbit anti-*Saccharomyces cerevisiae* PMA1 (sc-33735, Santa Cruz Biotechnology); rabbit anti-human CA-IX (ab15086, Abcam, Cambridge, MA); rabbit anti-human MCT1 (sc-50324, Santa Cruz, CA); rabbit anti-human GLUT1 (ab15309, Abcam, Cambridge, MA); rabbit anti-human NHE1 (sc-28758, Santa Cruz, CA); rabbit anti-human ER(#RM9101,ThermoFisher Scientific, MA). Histological stained slides were scanned using the Aperio ScanScope XT digital slide scanner and positivity analysis for each target gene staining was carried out using Aperio ImageScope V 10.2.1.2314 software. Positive cell percentage was calculated for PMA1, CA9, MCT1, GLUT1, and NHE1 expression on the entire tissue cross-section using algorithm membrane 9 in which positive cells include the cells with (3_+_) strong; (2_+_) medium; (1_+_) weak membrane intensity staining.

### RNA analyses in formalin-fixed, paraffin-embedded (FFPE) tissue

FFPE tissue samples were cut in 10µm-thick sections on a microtome, and deparaffinized by deparaffinization solution (Qiagen, 19093). Total RNA was extracted from deparaffinized FFPE sections with the miRNeasy FFPE kit (Qiagen, 217504) following the manufacturer’s protocol. Real-time qRT-PCR analyses for PMA1 mRNA were described above in the qRT-PCR section. Experimental C_t_ values from PMA1 amplification were normalized with GAPDH C_t_ values and were expressed relative to MCF-7/MOCK-1 control C_t_ values.

### Statistical analyses

A two-tailed unpaired student T-test or Welch’s T-test was employed to determine statistical significance. Ordinary one-way ANOVA with Geisser Greenhouse correction and Tukey’s Multiple Comparison test, with a single pooled variance. A p-value of less than 0.05 was considered statistically significant or otherwise indicated. Kaplan Meier Curve was used to analyze overall survival in mouse models with Log-Rank Test curve comparison.

## Data and code availability

The authors declare that the data supporting the findings of this study are available within the paper and its Supplementary information files. The code is available in the Supplementary data. Figures with raw data associated include Fig. 5H, associated raw data are found in Supplementary figures S19.

## Acknowledgements

**NIH U54 CA193489 (RJG); NIH R01 CA 077575-17 (RJG); NIH** F99 **CA234942** (**SR**); NIH P30 CA076292 (**core grant**)**; ERC Consolidator Award SURVIVE #723997 (PS)**

## Appendix

**Supplementary Table 1:** Average intracellular pHi in control and CA-IX expressing MCF-7 cells as a function of extracellular pH. n=158-438 cells , standard deviation shown.

| pHe | pHi |  |  |  |  |  |
| --- | --- | --- | --- | --- | --- | --- |
|  | MCF7 |  | M1 |  | M6 |  |
|  | Average | SD | Average | SD | Average | SD |
| 6.4 | 6.78 | 0.1966 | 6.78 | 0.178 | 6.75 | 0.1859 |
| 6.8 | 6.88 | 0.1459 | 6.93 | 0.1224 | 6.97 | 0.08992 |
| 7.1 | 6.98 | 0.1262 | 7.04 | 0.08474 | 7.04 | 0.09440 |
| 7.4 | 7.05 | 0.1517 | 7.04 | 0.09170 | 7.03 | 0.07948 |

**Supplementary Table 2:** Effect of PMA1 expression on experimental and spontaneous metastasis in MCF-7 cells. (Two-tailed Fisher’s exact t-test p<0.05*)

| Mouse Model | Experimental Metastasis | Spontaneous Metastasis |
| --- | --- | --- |
|  | # of mice with lung mets | # of mice with lung mets |
| Group ID |  |  |
| MCF7/PMA-1 | 7/12 (58%) | 4/9 (44%) |
| MCF7/MOCK-1 | 1/10 (10%) | 1/10 (10%) |
| P-Value | 0.03* | 0.14 (n.s.) |

**Supplementary Fig. 1.**
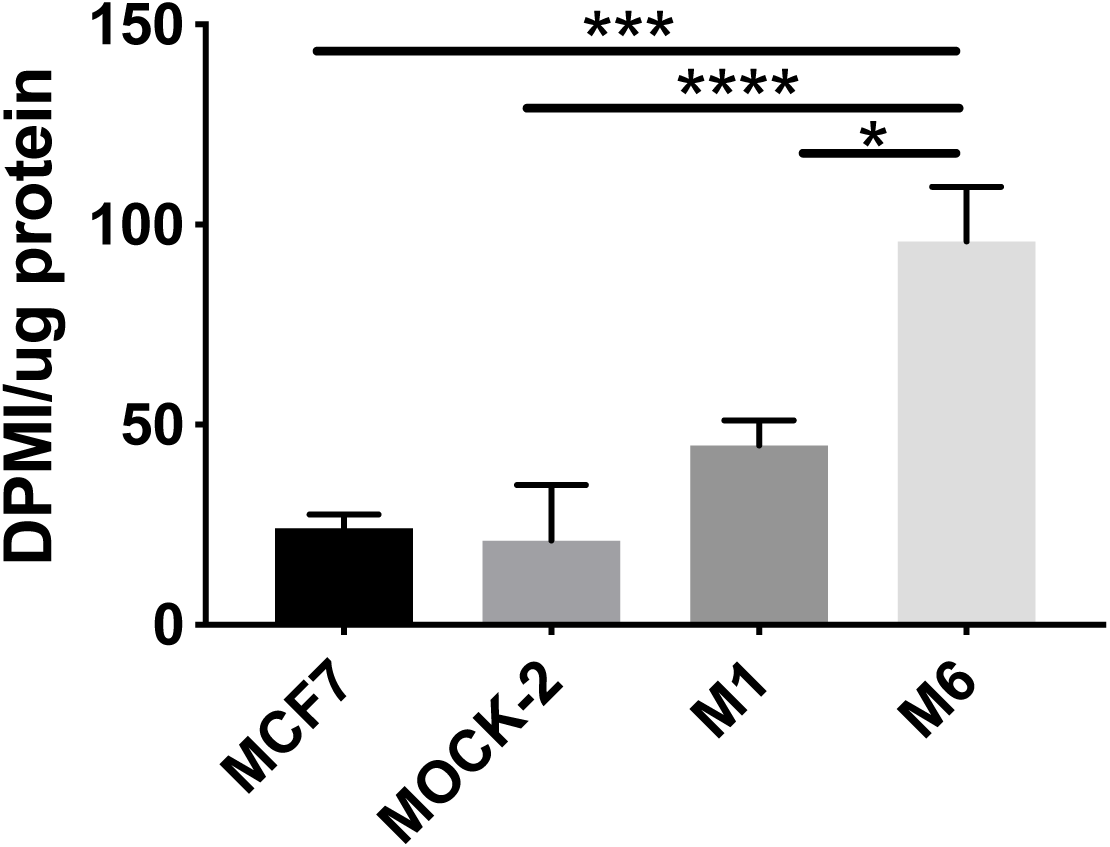
Uptake of ^3^H-2-deoxy glucose in control and CAIX MCF-7 clones (N=3; Ordinary one-way ANOVA *, ***, **** p<0.05, 0.005, 0.001)

**Supplementary Fig. 2.**
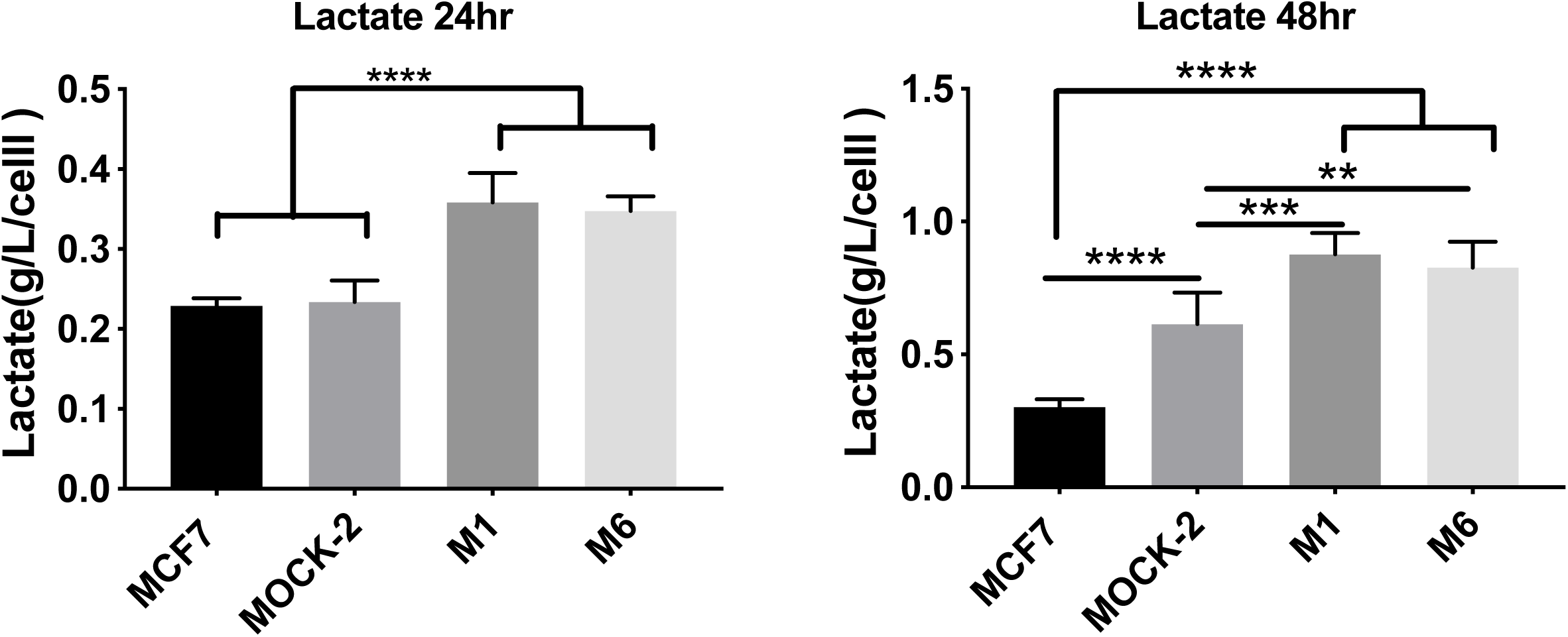
YSI analysis of lactate in media at 24 and 48 hours in CA-IX clones (N=6 replicates, Ordinary one-way ANOVA p<0.01**, 0.005***, 0.001****.)

**Supplementary Fig. 3.**
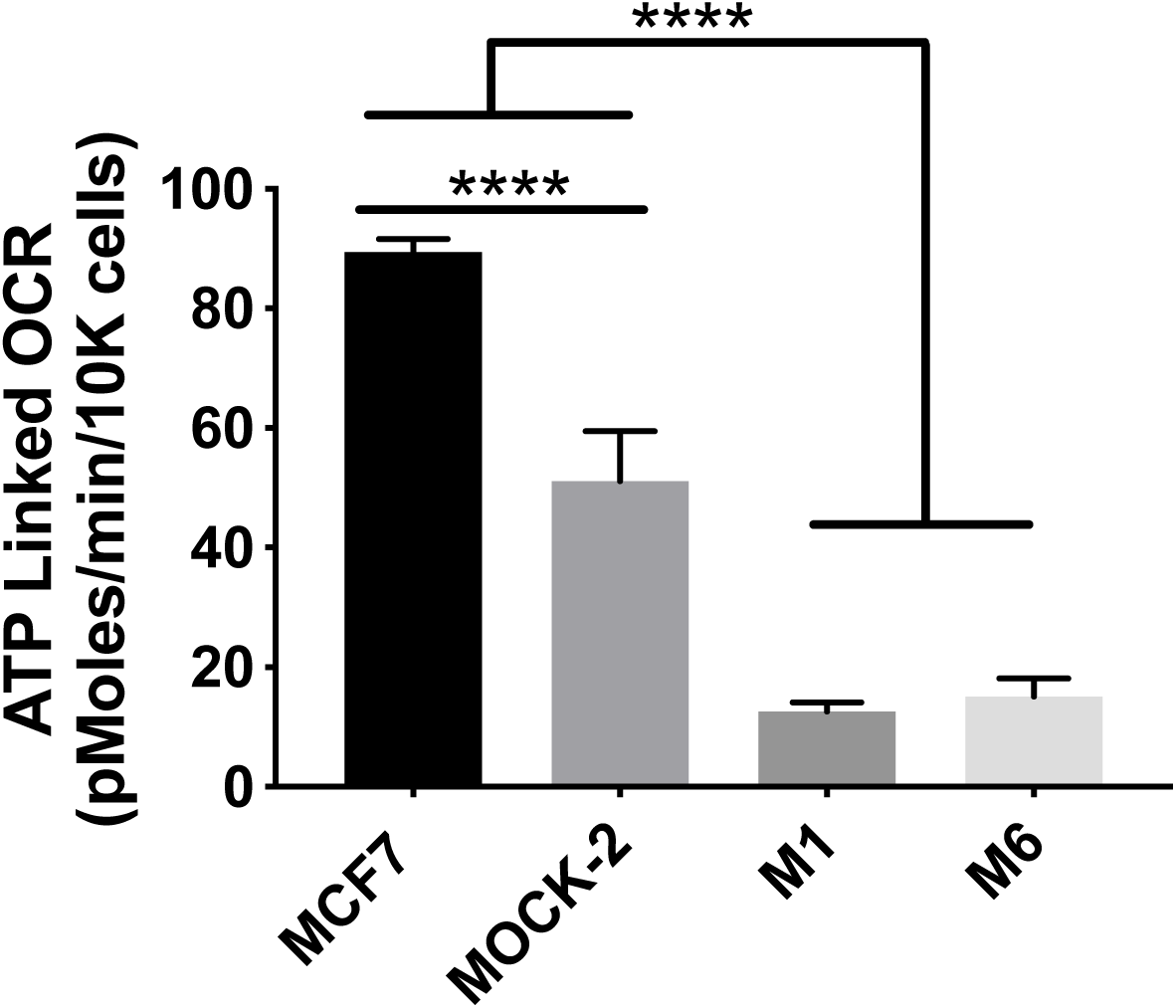
ATP linked OCR measured by mitochondrial stress test XFe96 Seahorse assay, by injecting oligomycin to shut off mitochondrial respiration. (N=8 replicates, ordinary one-way ANOVA p<0.001****.)

**Supplementary Fig. 4.**
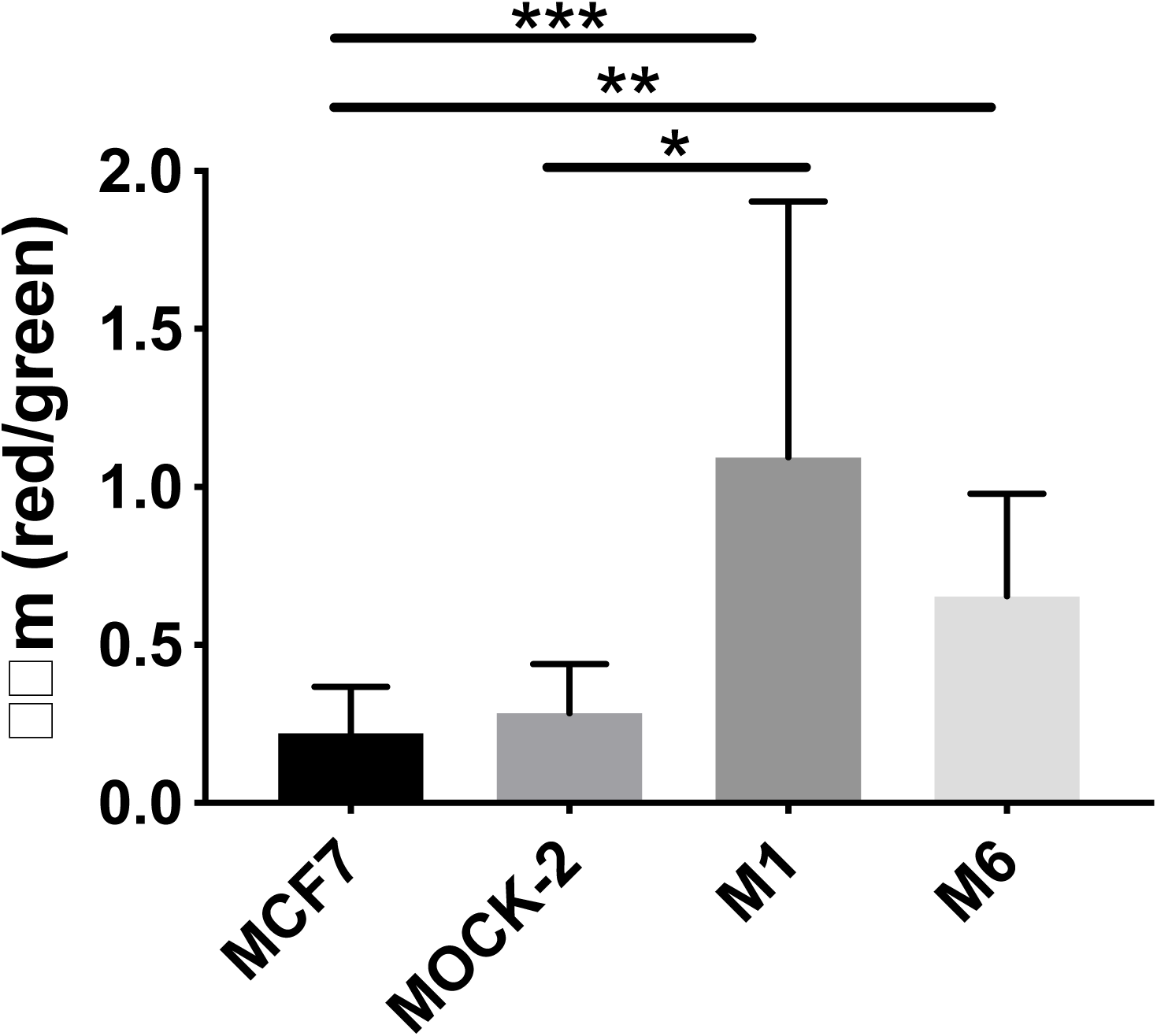
Mitochondrial polarization using JC-1 mitochondrial dye in CA-IX clones (N=8-12 per group, 2 bioreplicates; Kriskal Wallis Test p<0.05*,p<0.01**, p<0.005***)

**Supplementary Fig. 5.**
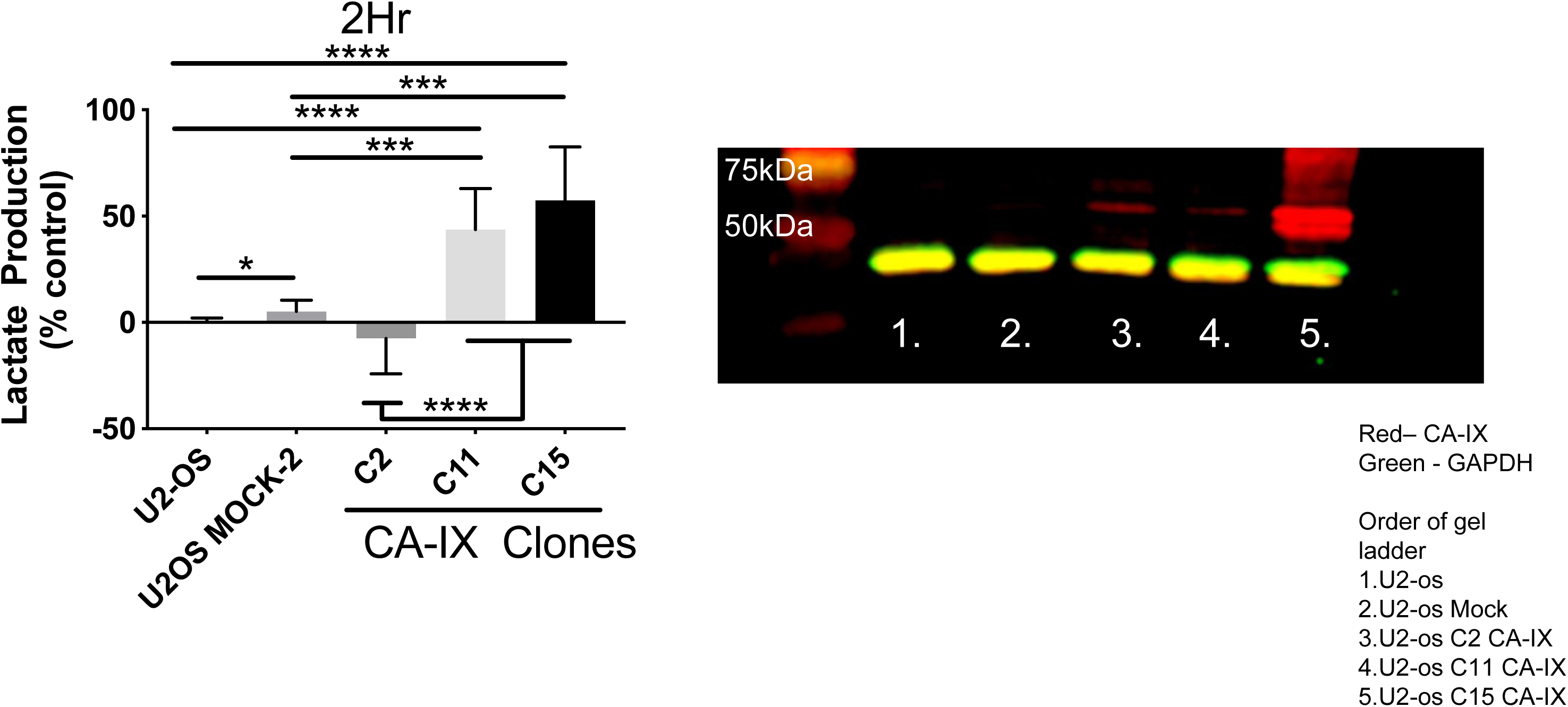
U2OS Lactate production rates and CA-IX western blot (N=12; Brown Forsythe and Welch ANOVA, p<0.05*, p<0.01**, p<0.005***, p<0.001****.)

**Supplementary Fig. 6.**
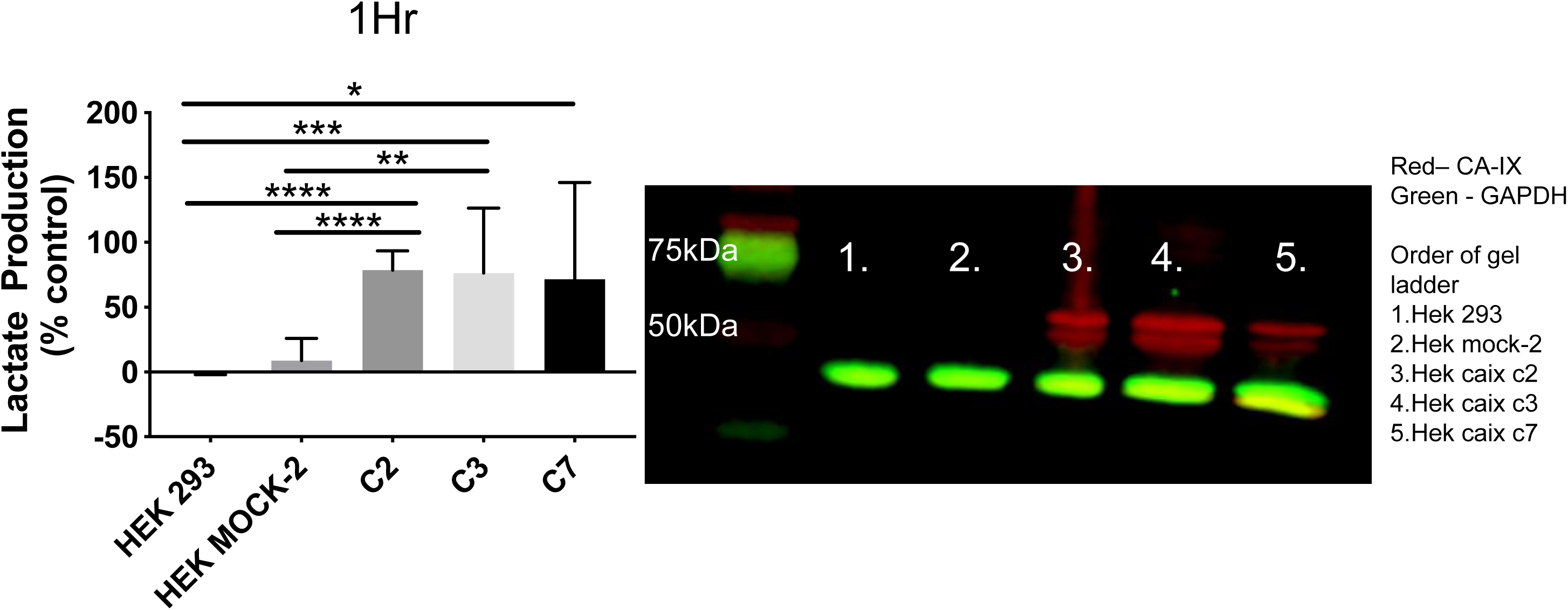
Lactate production and Western blot of CA-IX transfected HEK 293T cells. (N=15; Brown Forsythe and Welch ANOVA, p<0.05*, p<0.01**, p<0.005***, p<0.001****.)

**Supplementary Fig 7.**
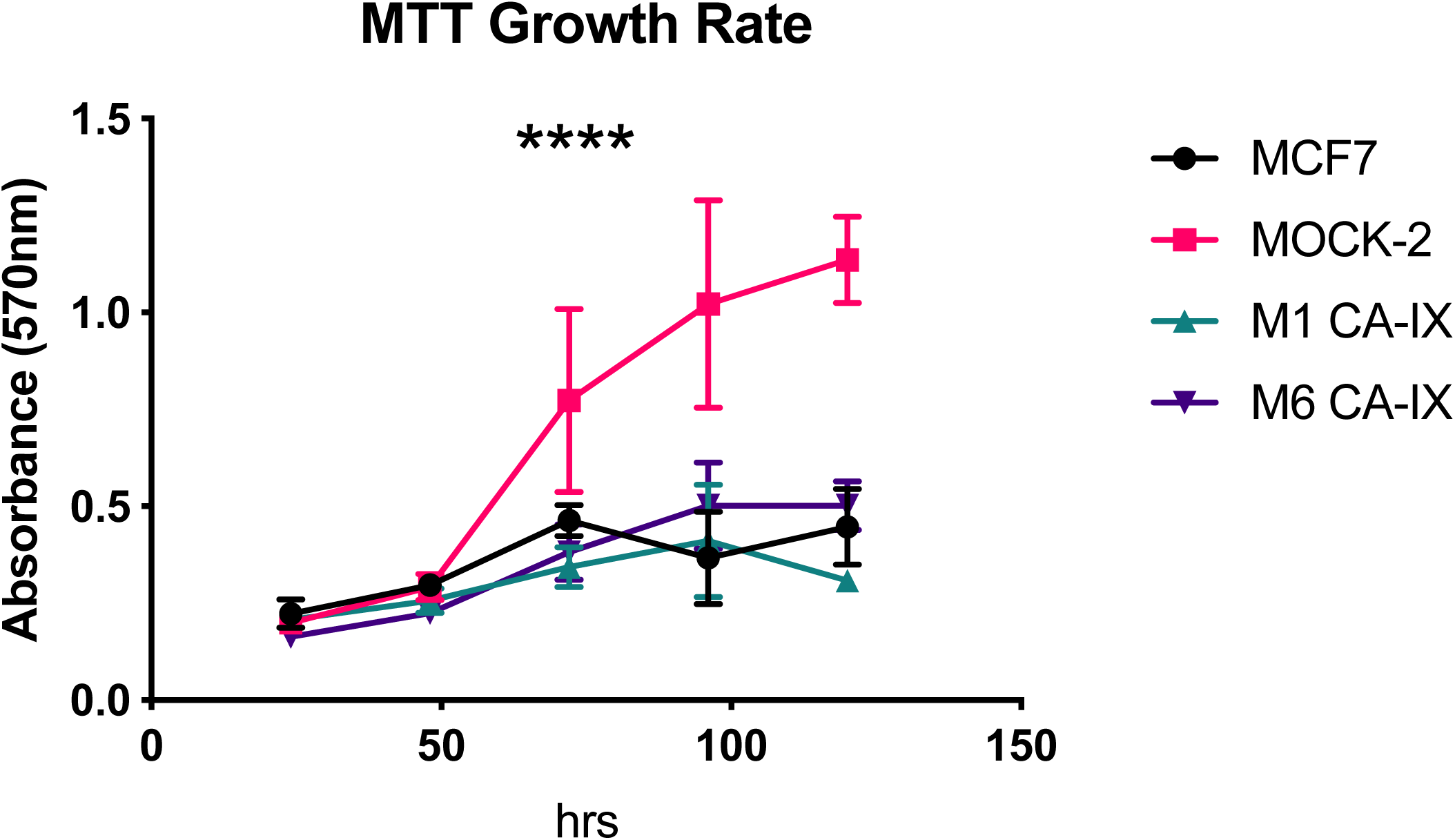
In vitro growth rates of MCF-7, MOCK-2 and CA-IX clones M1 and M6. n= 3, average ± SD. p<0.0001****.

**Supplementary Fig. 8.**
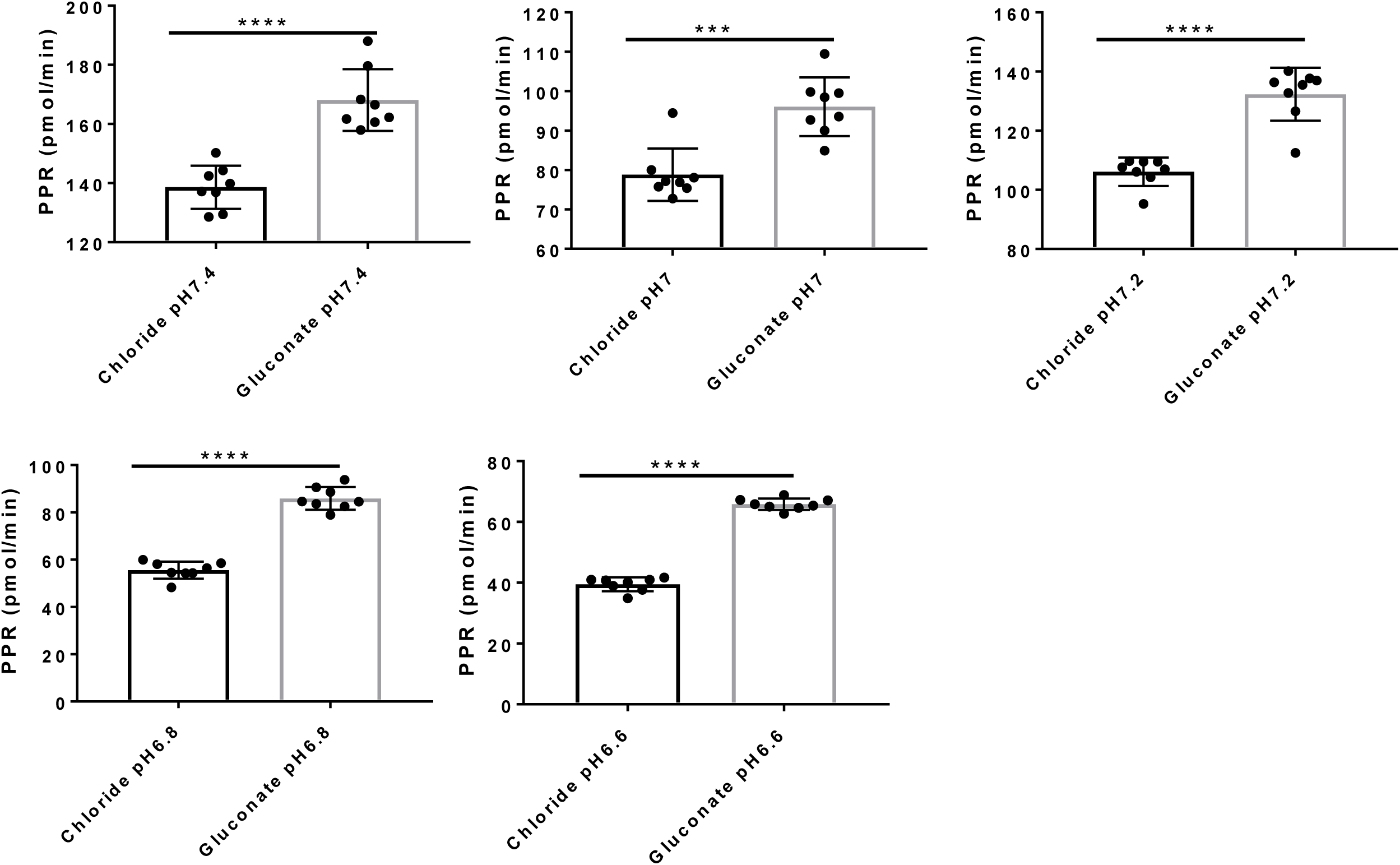
Effect of chloride vs gluconate on PPR. Average PPR ± SD, n=8, unpaired t-test p<0.005***, p<0.0001****.

**Supplementary Fig. 9.**
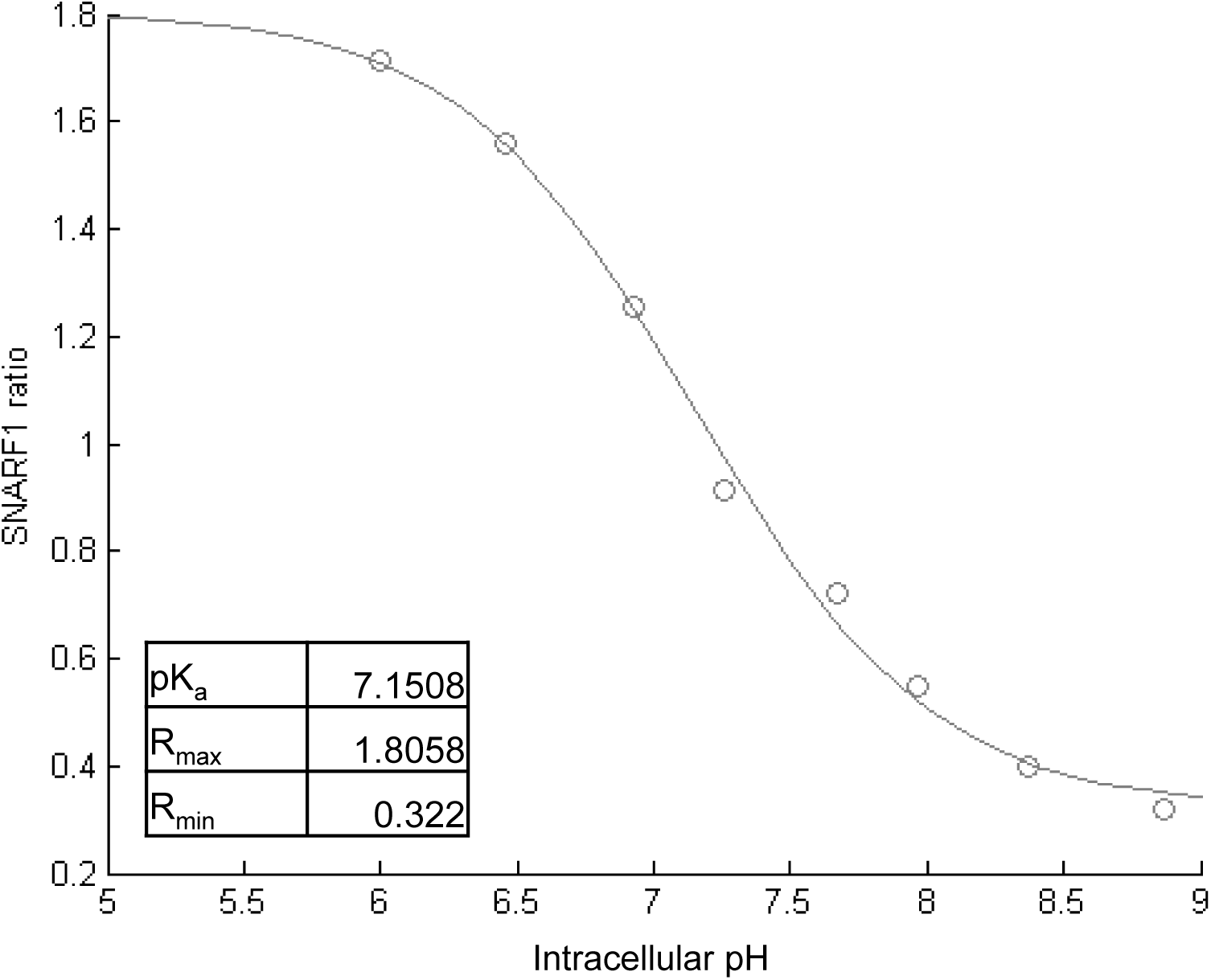
cSNARF1 calibration curve with nigericin/high K^+^

**Supplementary Fig. 10.**
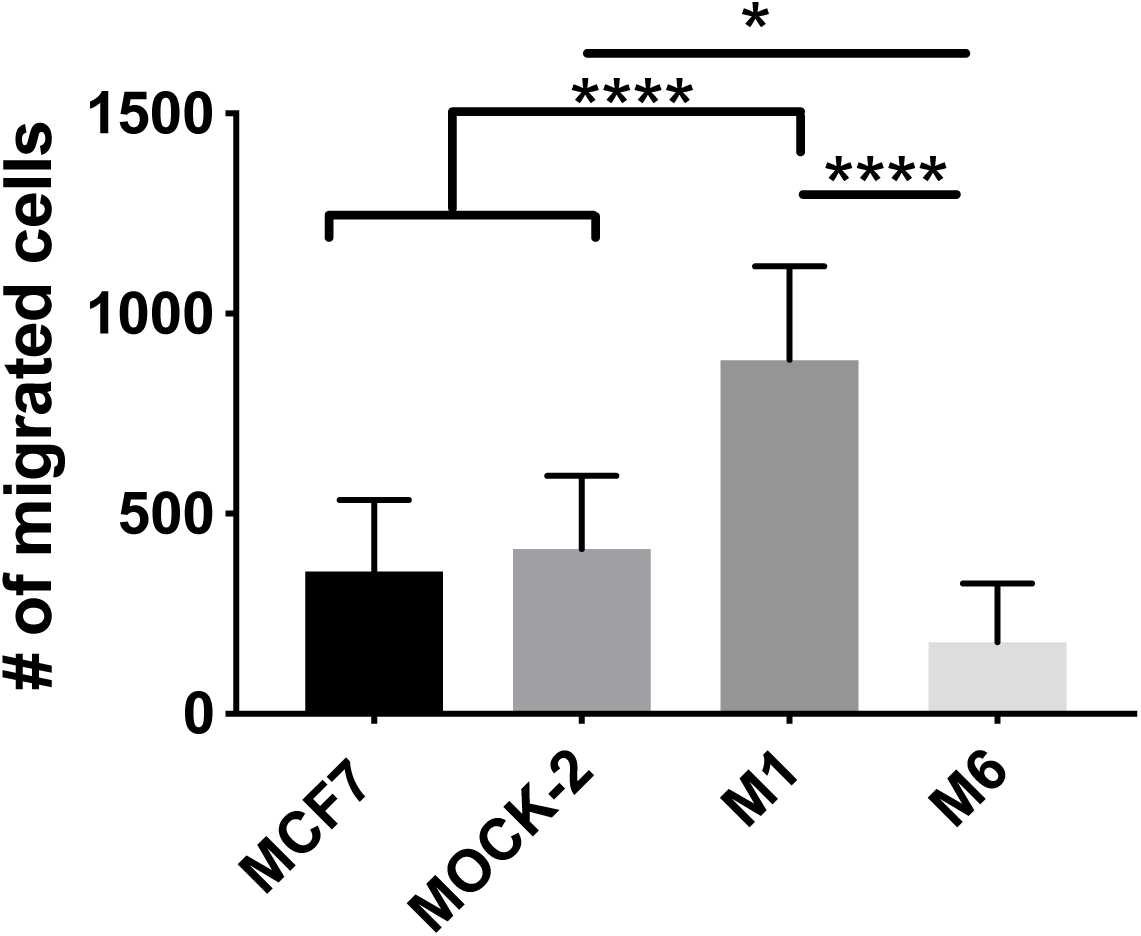
Migration assay to measure effects of CA-IX expression on migration of MCF7 cells. Imaged using Celigo, n=3 bio-replicates in triplicate, average ± SD. (Ordinary one-way ANOVA, p<0.05*, p<0.005***,p<0.0001****)

**Supplementary Fig. 11.**
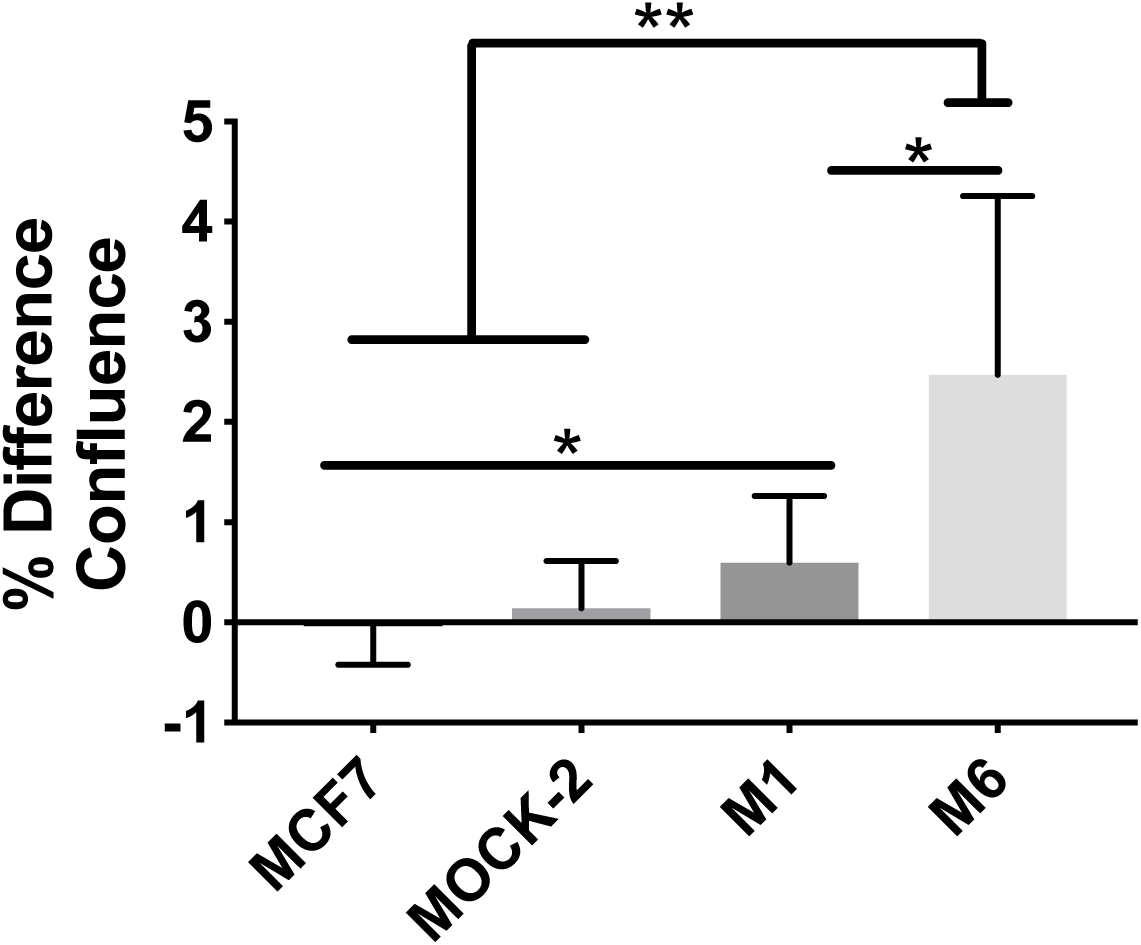
Gel escape assay to measure effects of CA-IX expression on invasion and migration in MCF7 cells. Imaged using Celigo, n=3 bio-replicates in triplicate, average ± SD. (Brown-Forsythe and Welch ANOVA, p<0.0001****)

**Supplementary Fig. 12.**
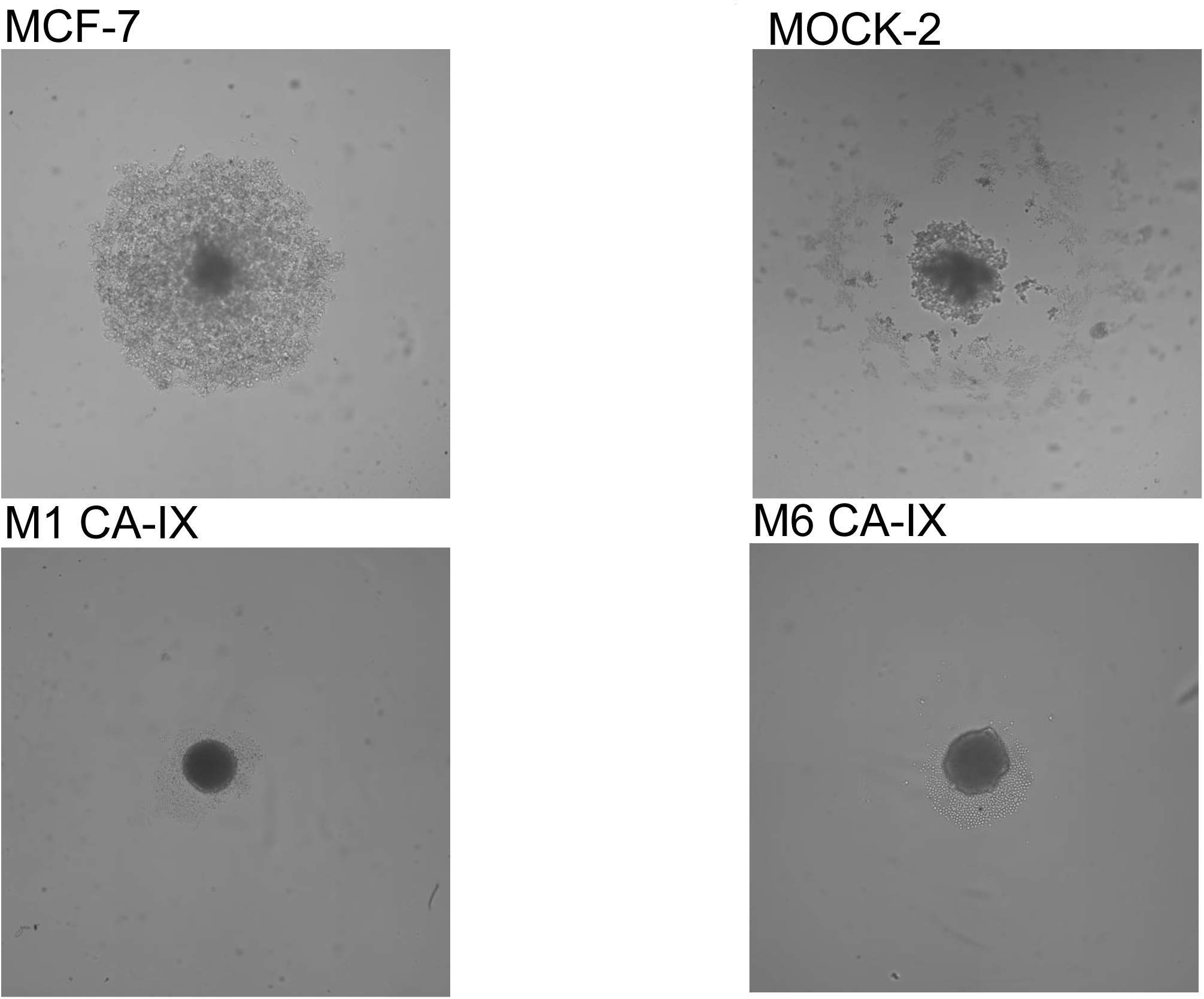
Spheroid formation assay to measure ability of cells to grow in 3D, an indicator of metastatic ability *in vitro*. Imaged using Celigo with single colony verification analysis after 5 days of growth in hanging drop plates. Representative images shown.

**Supplementary Figure 13:**
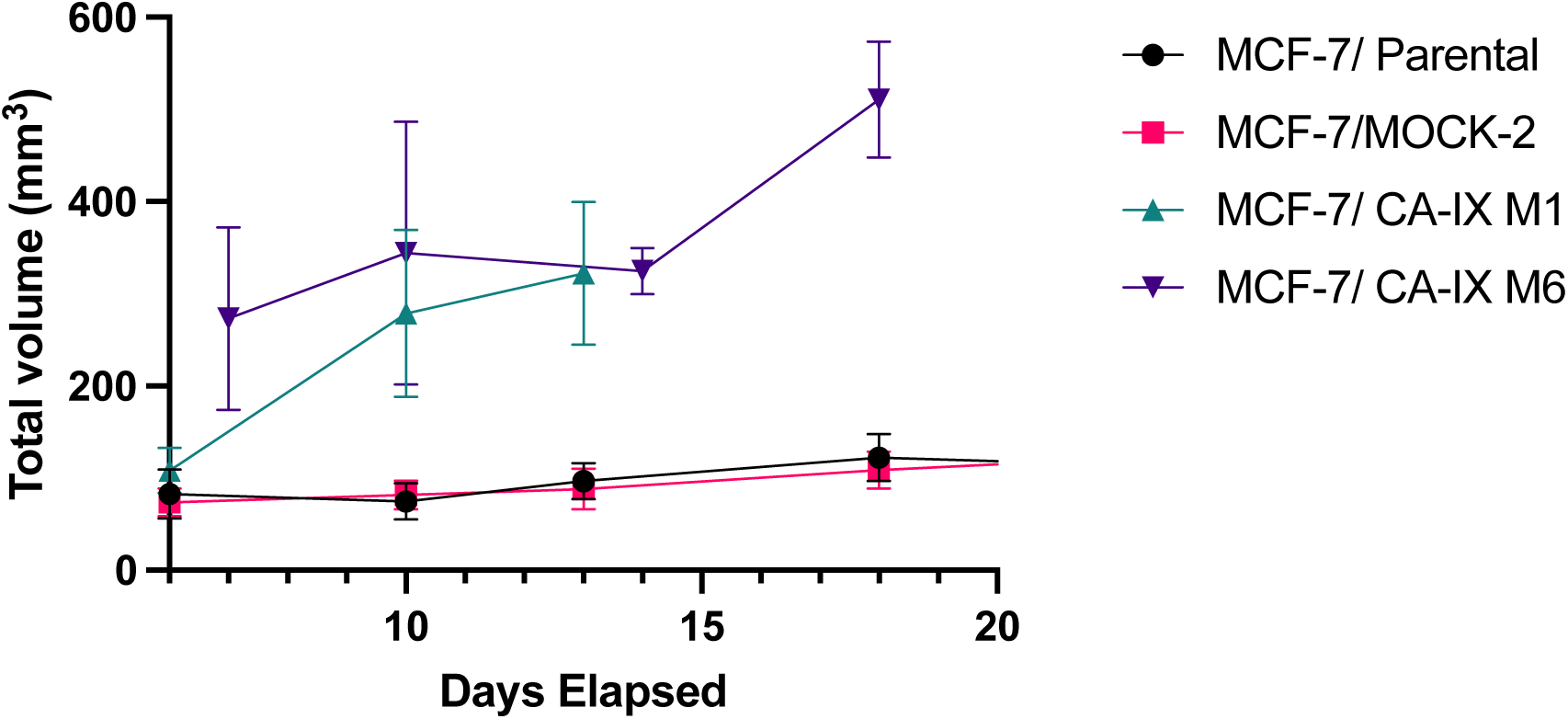
Effect of CA-IX expression on primary tumor growth of MCF-7 cells. n=12-15 mice per group

**Supplementary Fig. 14.**
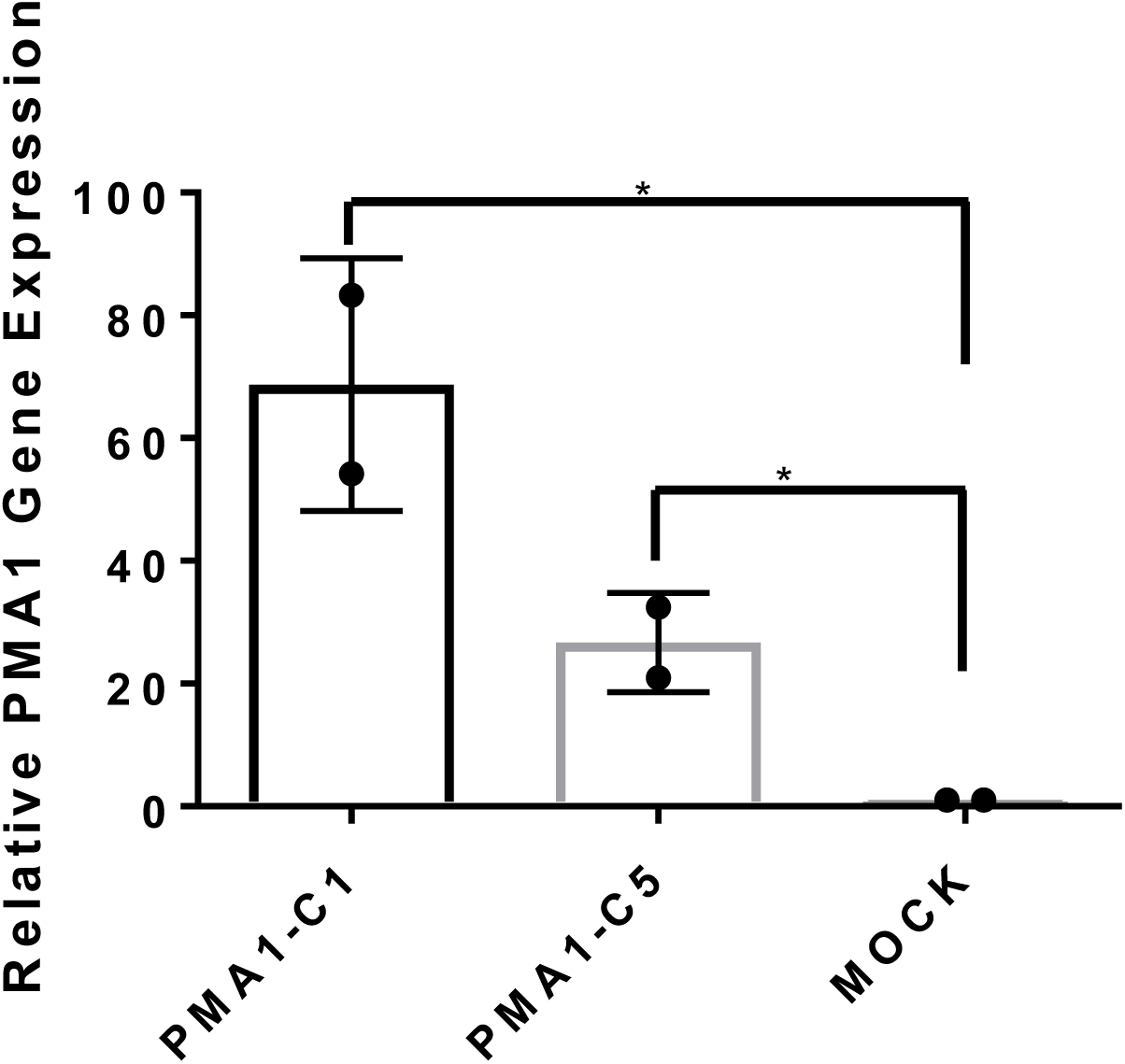
qRT PCR of relative PMA1 gene expression in PMA1 transfected clones compared to MOCK (empty vector clone). Average ± SD. (Ordinary one-way ANOVA. p<0.05*)

**Supplementary Fig. 15.**
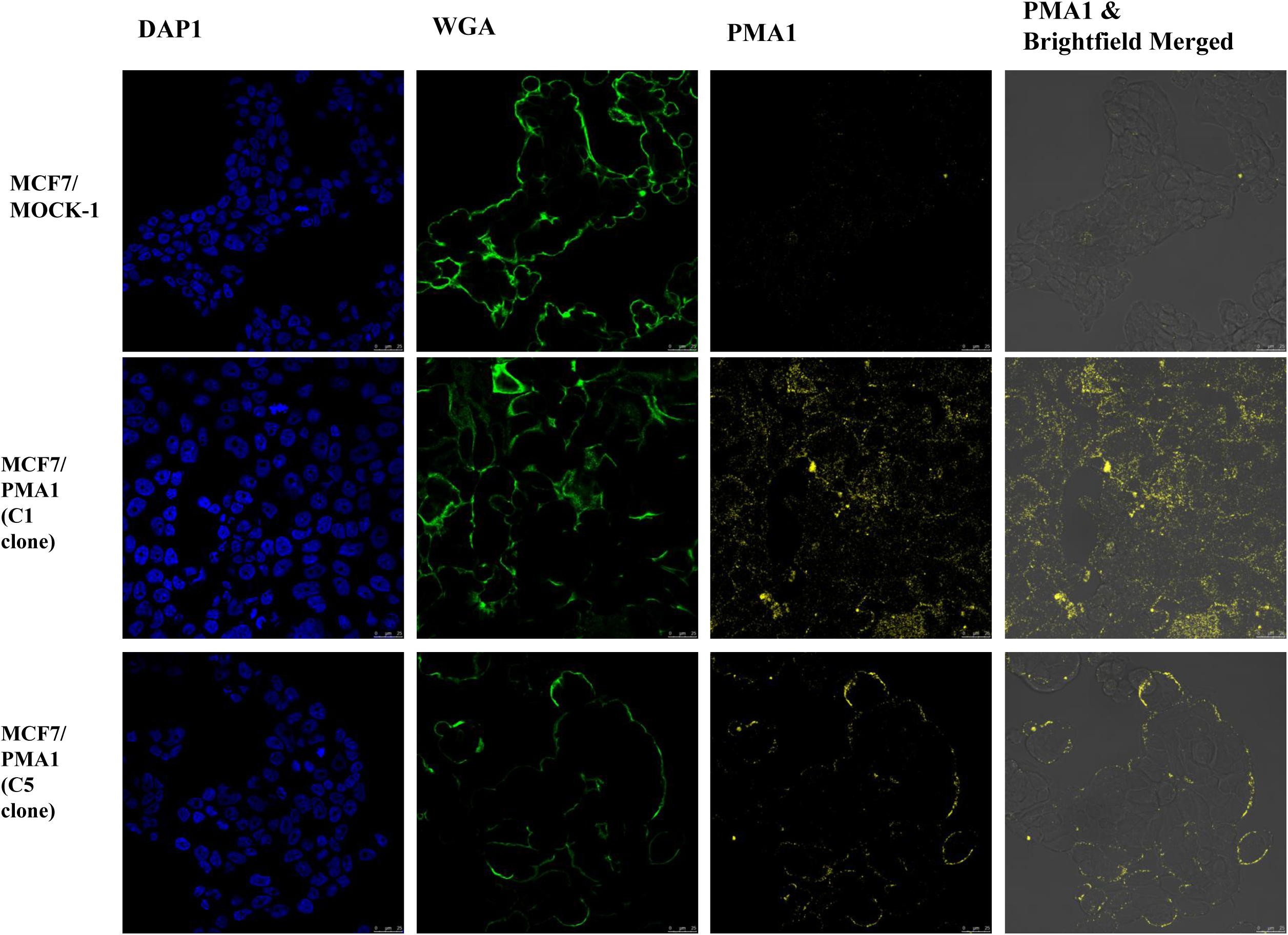
Characterization of PMA1-expressing cells by ICC staining (non-permeabilized) DAPI (blue), WGA (green), PMA1 (Alexa Fluor 594 red)

**Supplementary Fig. 16.**
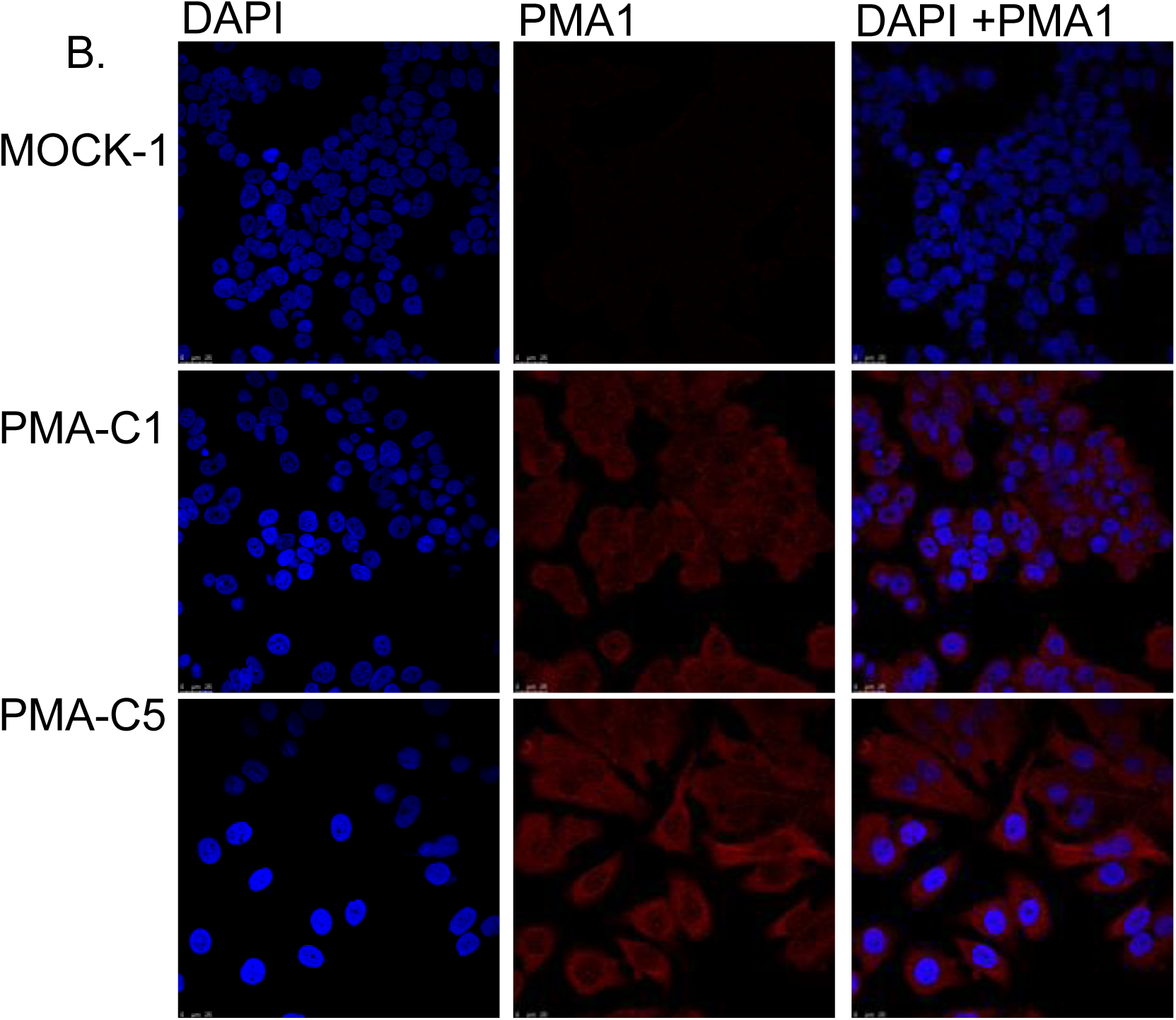
ICC of permeabilized cells staining for PMA1. PMA in red and Nuclear stain DAPI in blue.

**Supplementary Fig. 17.**
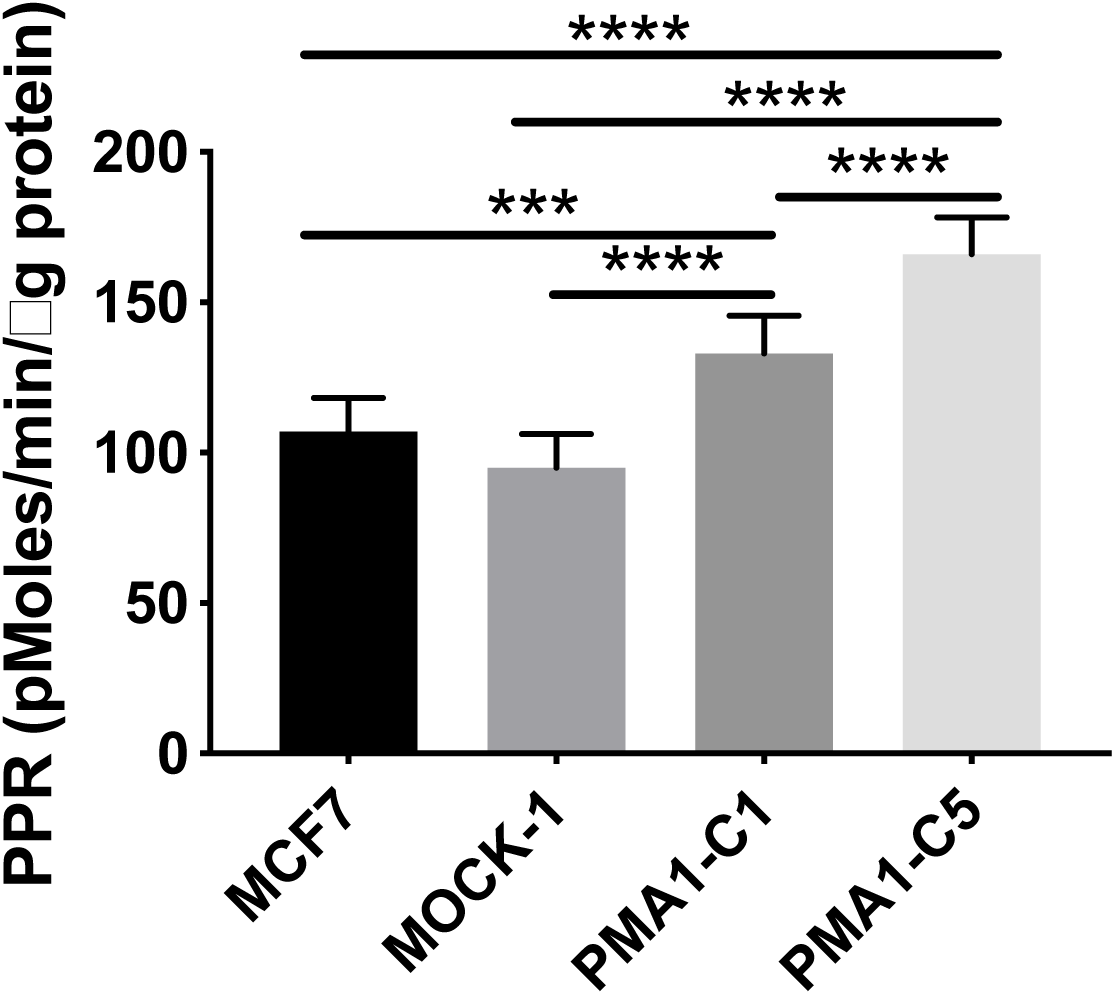
Glycolytic reserve measured by glycolysis stress test with XFe96 Seahorse assay. Reserve is determined by difference in glucose-stimulated proton production rate (PPR) and addition of oligomycin which inhibits mitochondrial energy production. Average ± SD. (Ordinary one-way ANOVA. p<0.0005***, p<0.0001****)

**Supplementary Fig. 18.**
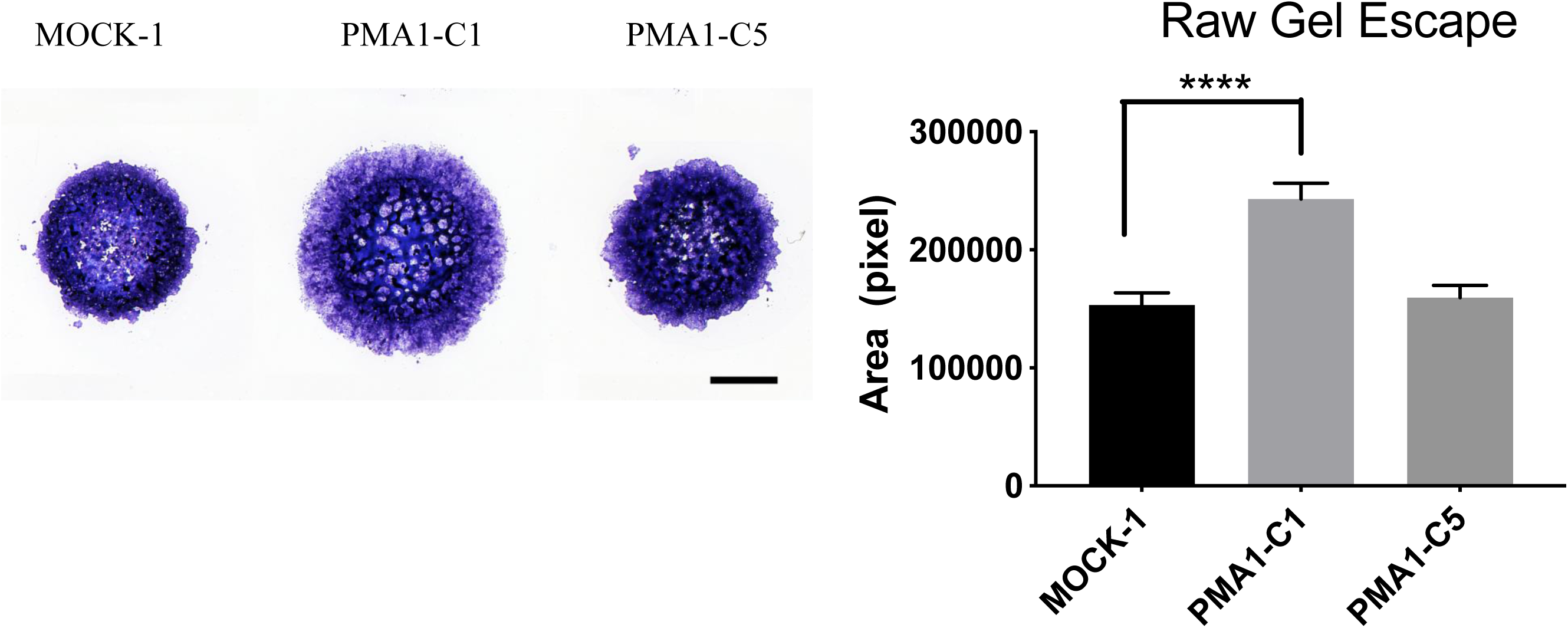
Raw data of gel escape assay and corresponding images, prior to normalization by growth rate. N=4 biological replicates, average ± SD. (Ordinary one-way ANOVA, p<0.0001****)

**Supplementary Fig. 19.**
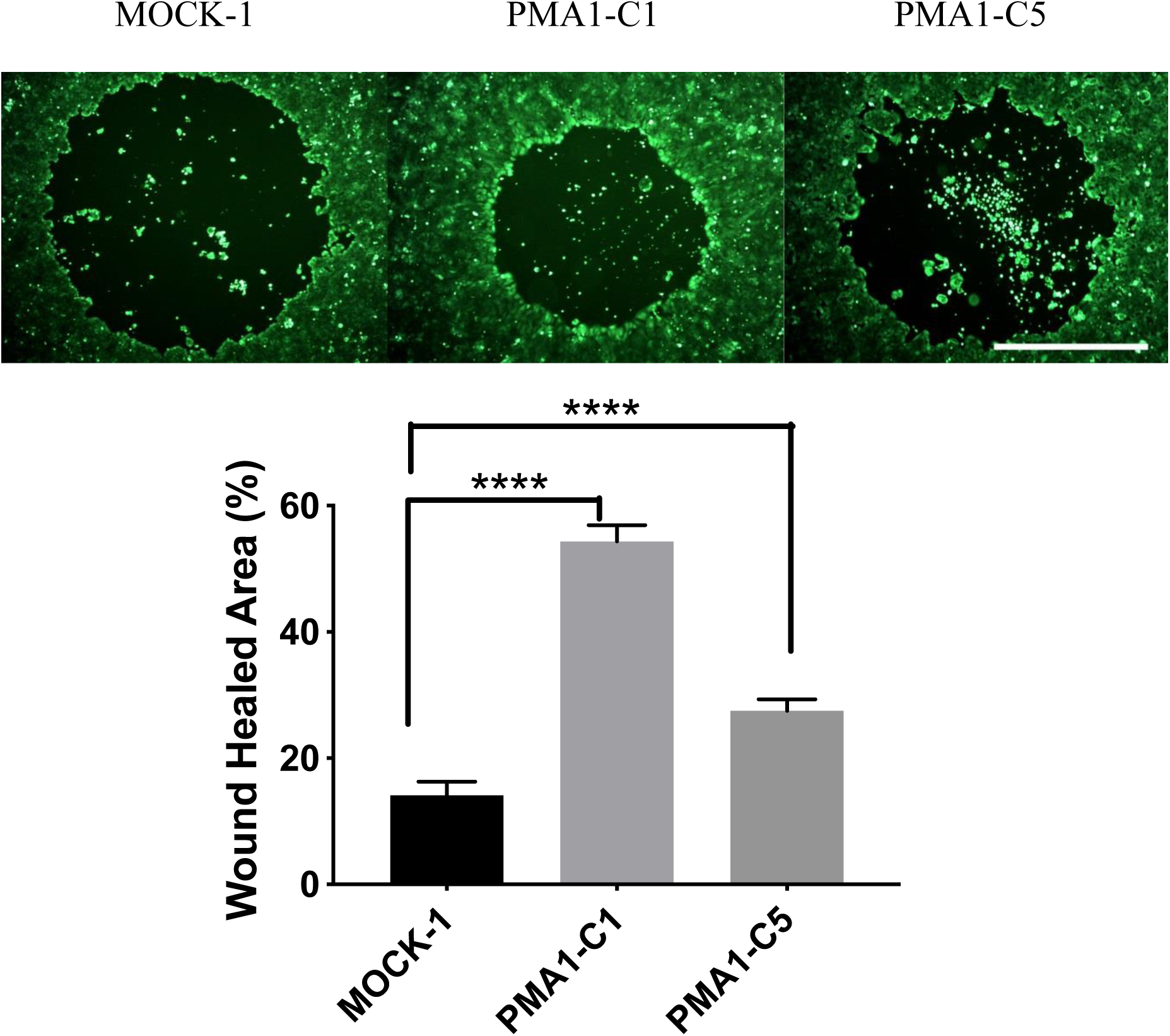
Raw data of circular wound healing assay and corresponding images, prior to normalization by growth rate. N=4 biological replicates, average ± SD. (Ordinary one-way ANOVA, p<0.0001****)

**Supplementary Fig. 20.**
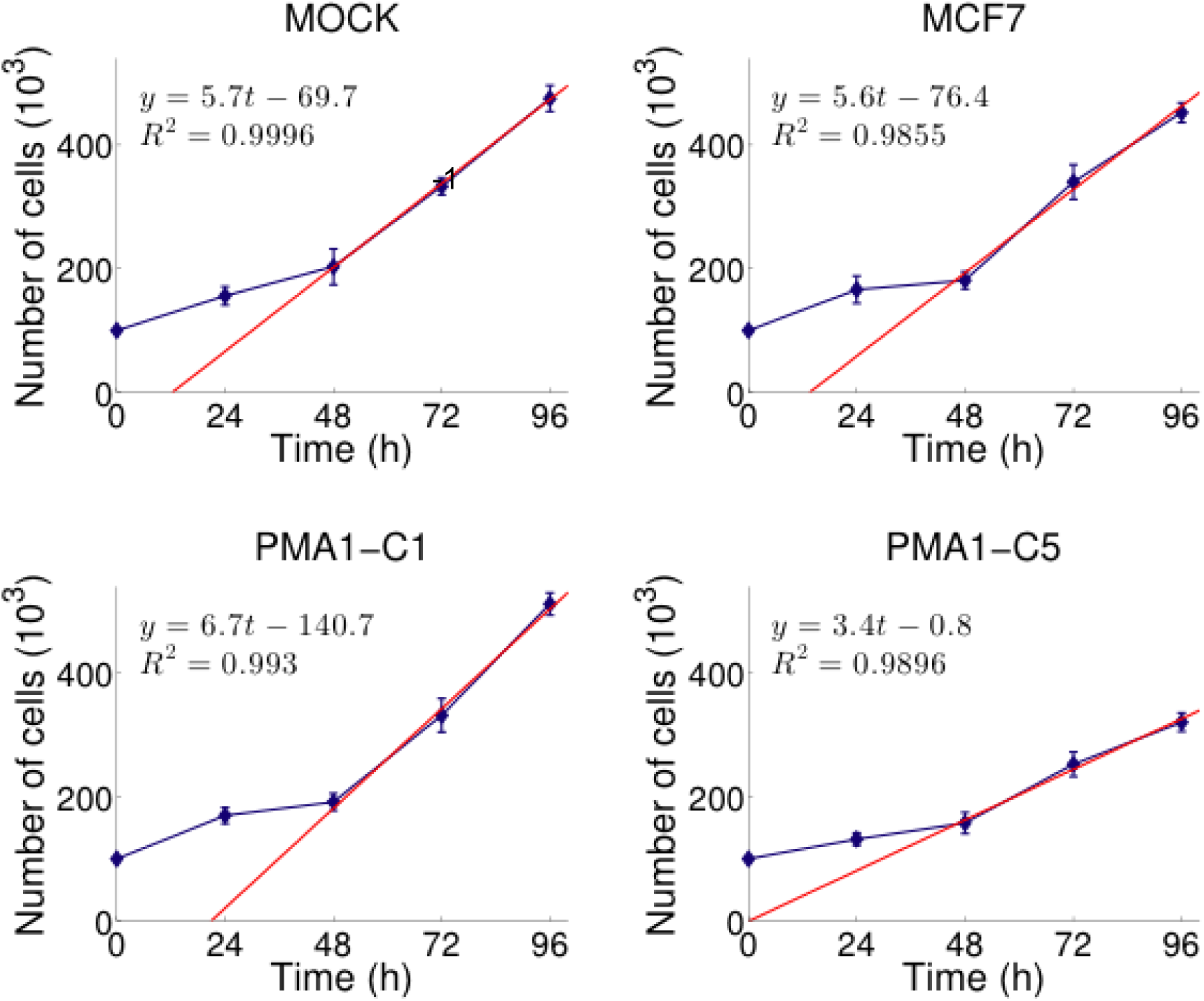
Cellular growth rate was determined by number of cells over time and calculated by a linear fit of cell growth for normalization of invasion and migration assays

**Supplementary Fig. 21.**
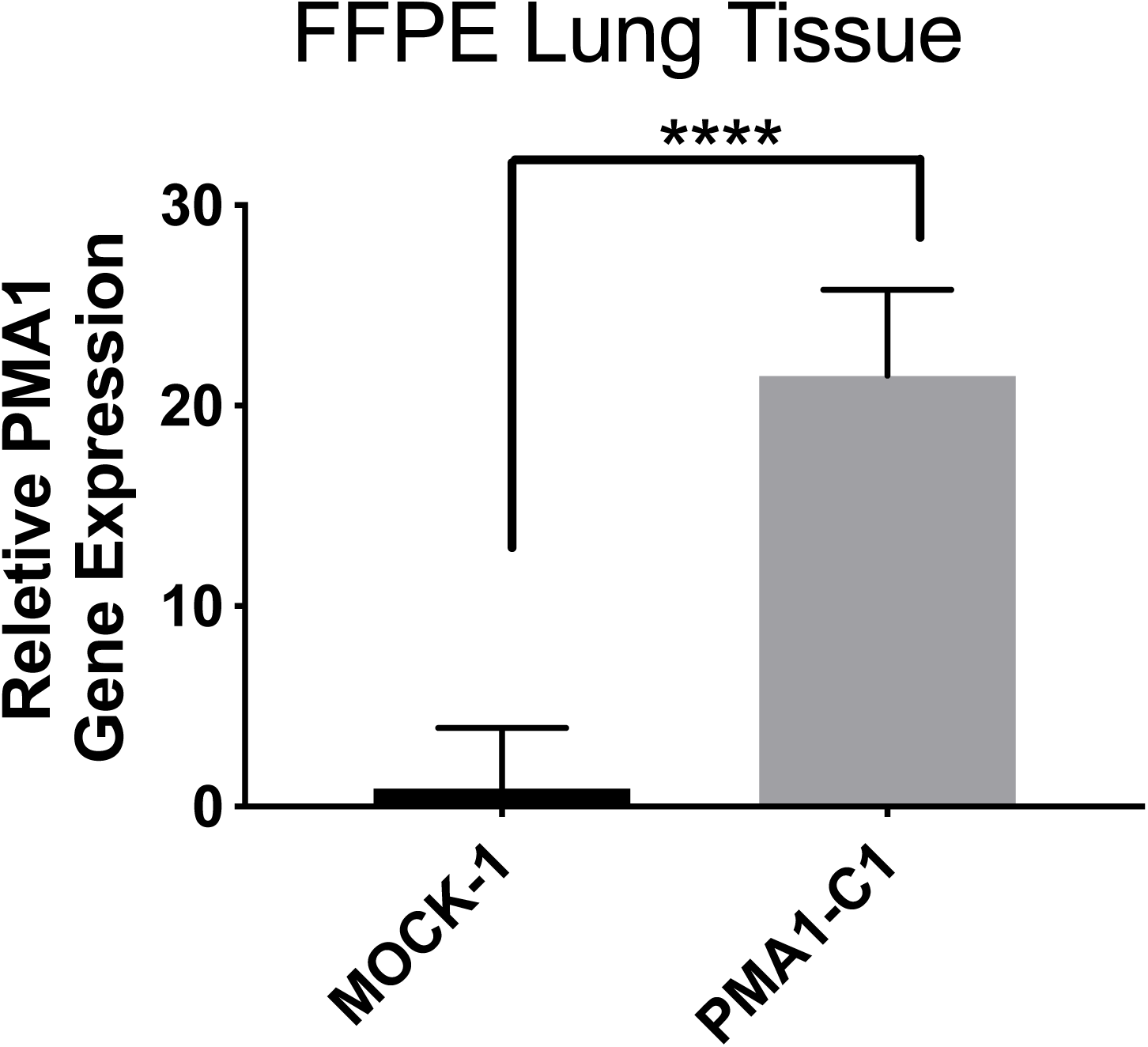
Relative PMA1 gene expression in lung micrometatases. Average relative gene expression ± SD, unpaired t-test p<0.0001.

**Supplementary Fig. 22.**
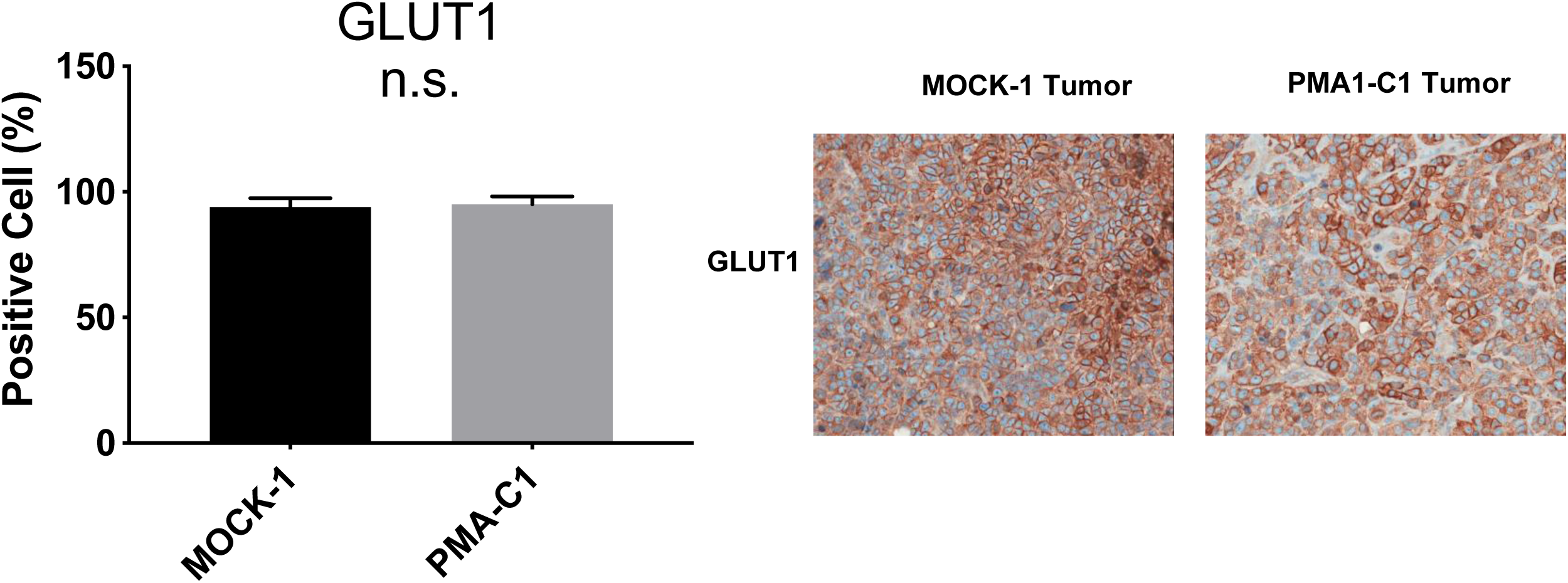
Quantification of GLUT1 protein staining in IHC samples of resected primary tumors and representative images. N=9-10, Average ± SD, unpaired t-test n.s.

**Supplementary Fig. 23.**
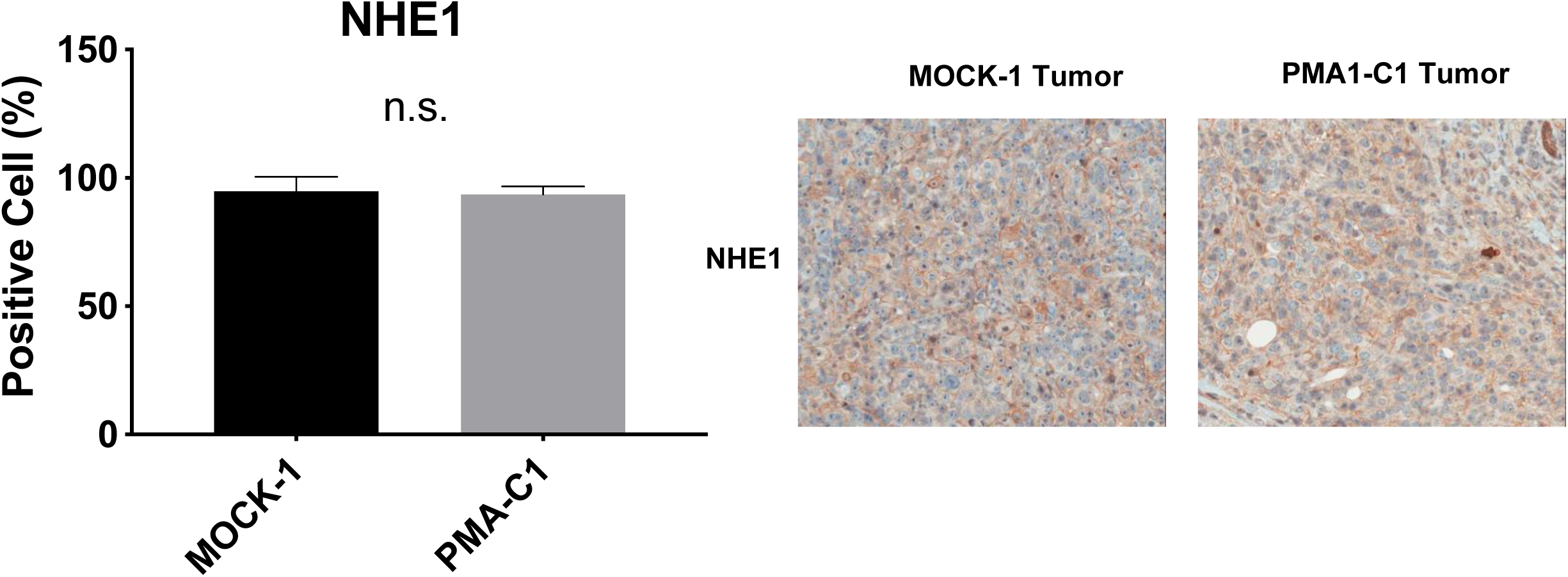
Quantification of NHE1 protein staining in IHC samples of resected primary tumors and representative images, n=8-9 Average ± SD, unpaired t-test n.s.

**Supplementary Fig. 24.**
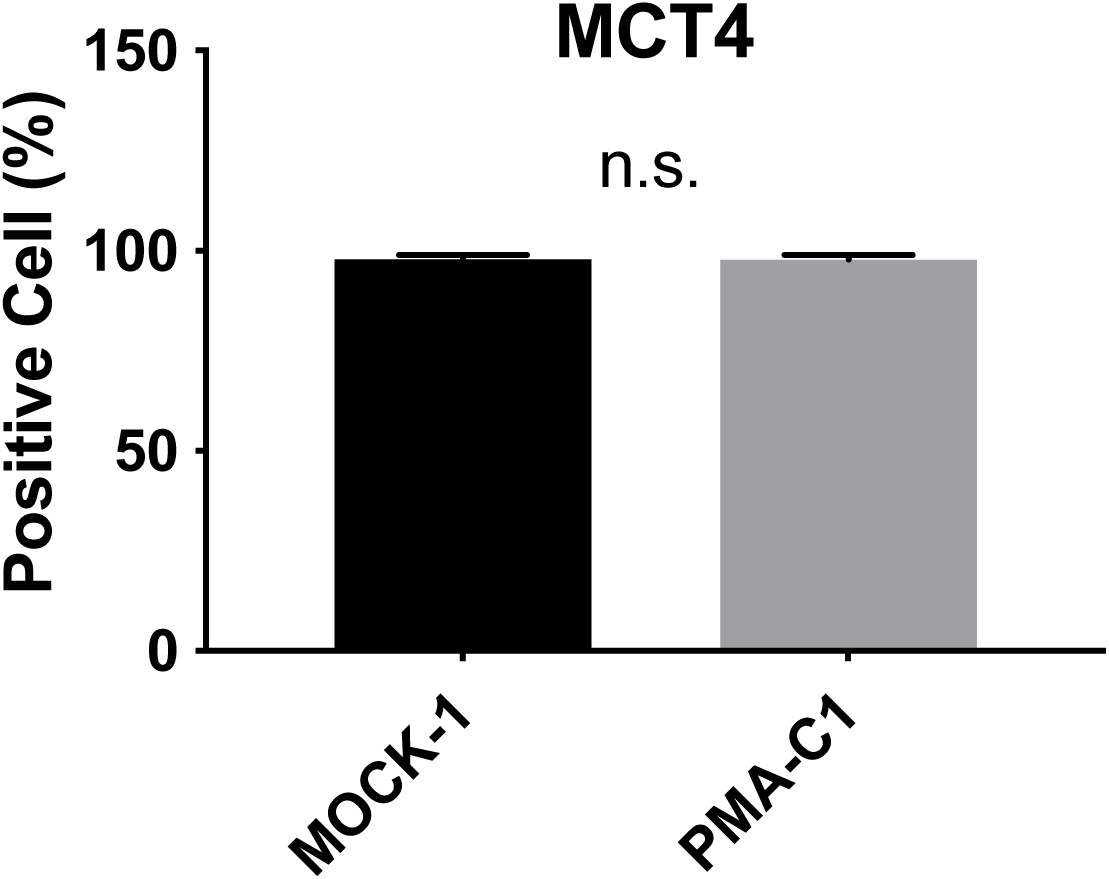
Quantification of MCT4 protein staining in IHC samples of resected primary tumors and representative images, n=8-9 Average ± SD, unpaired t-test n.s.

### Raw western blots

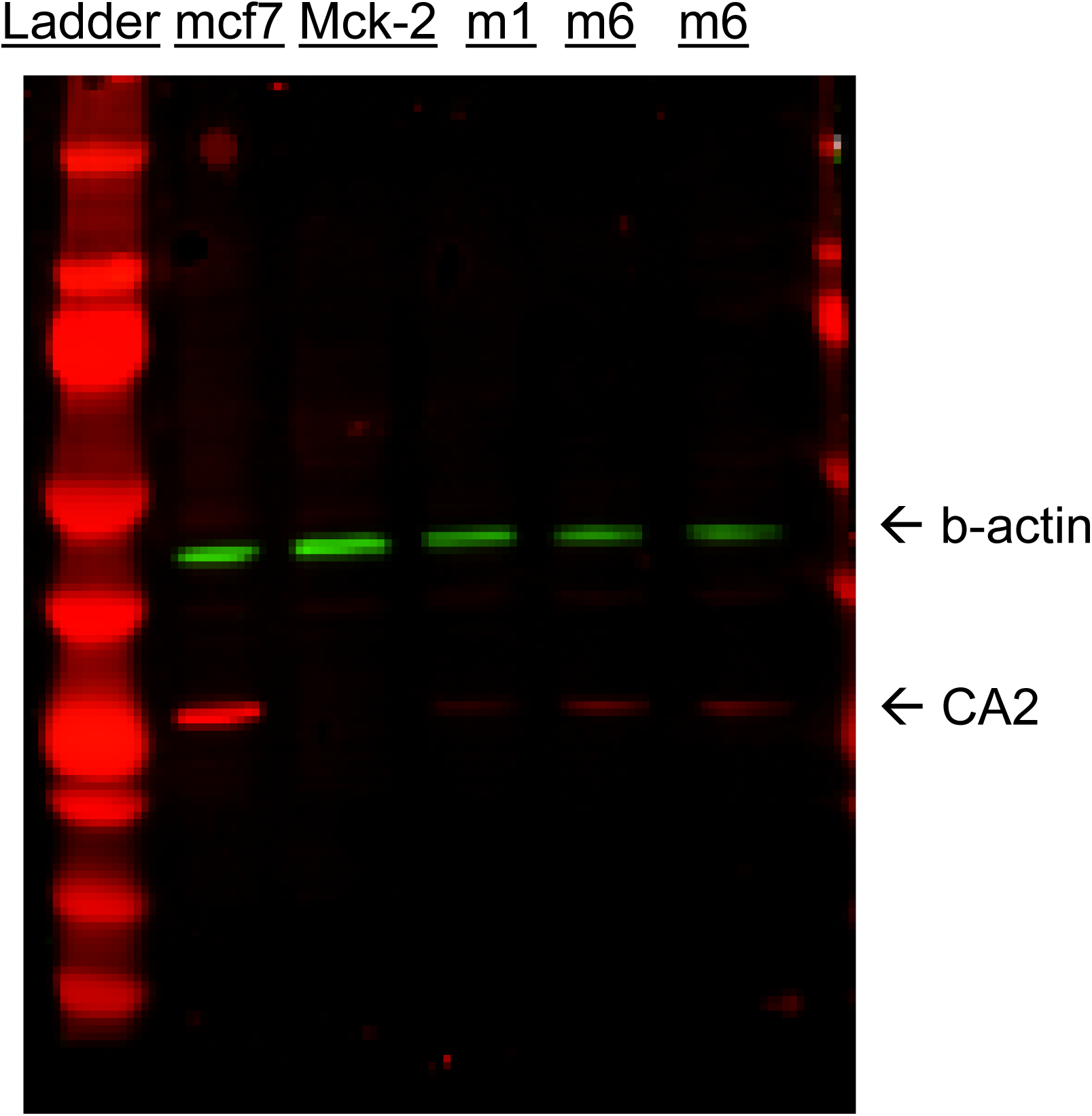

### Raw western

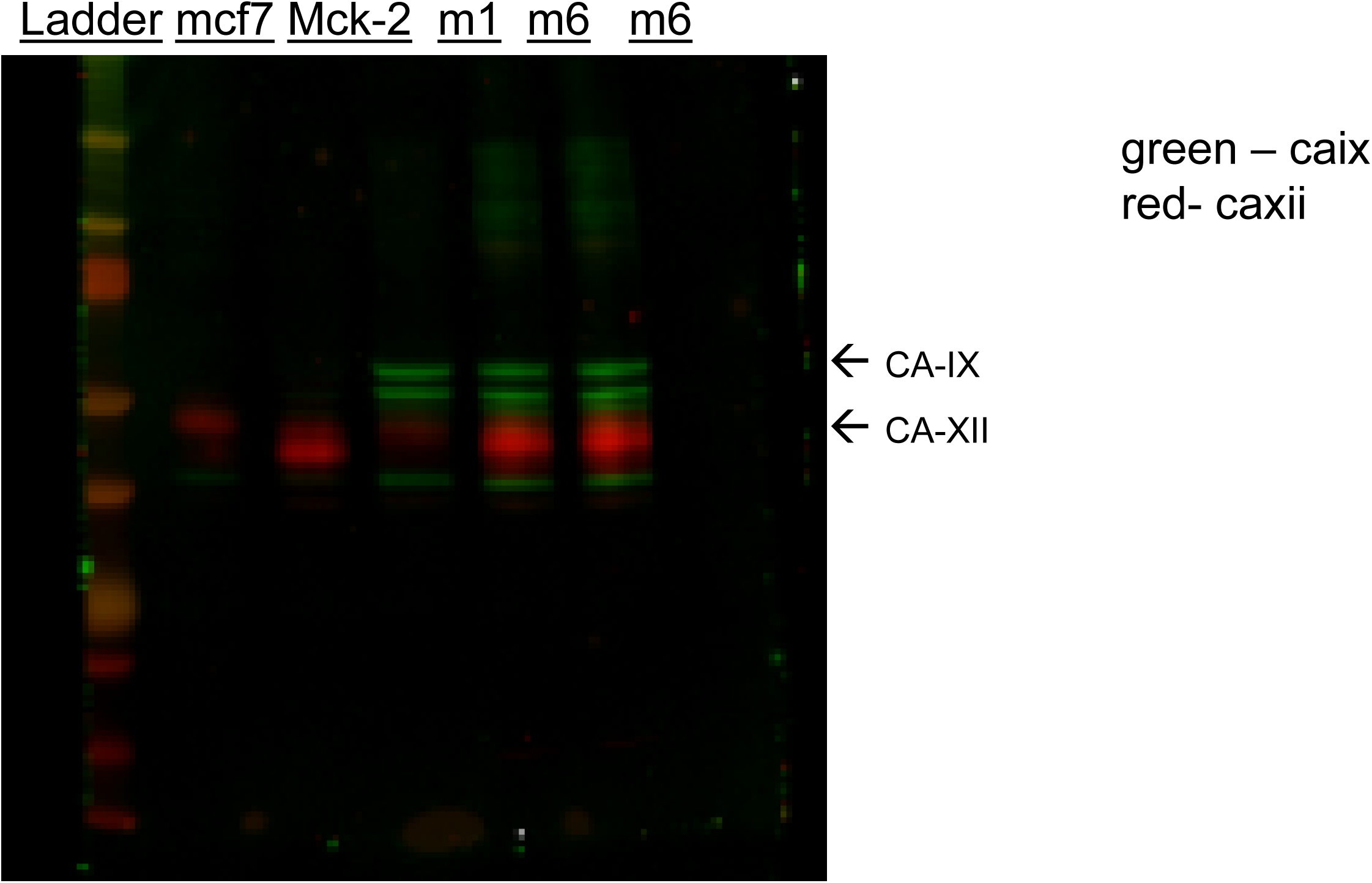

### Raw western u2OS CA-IX

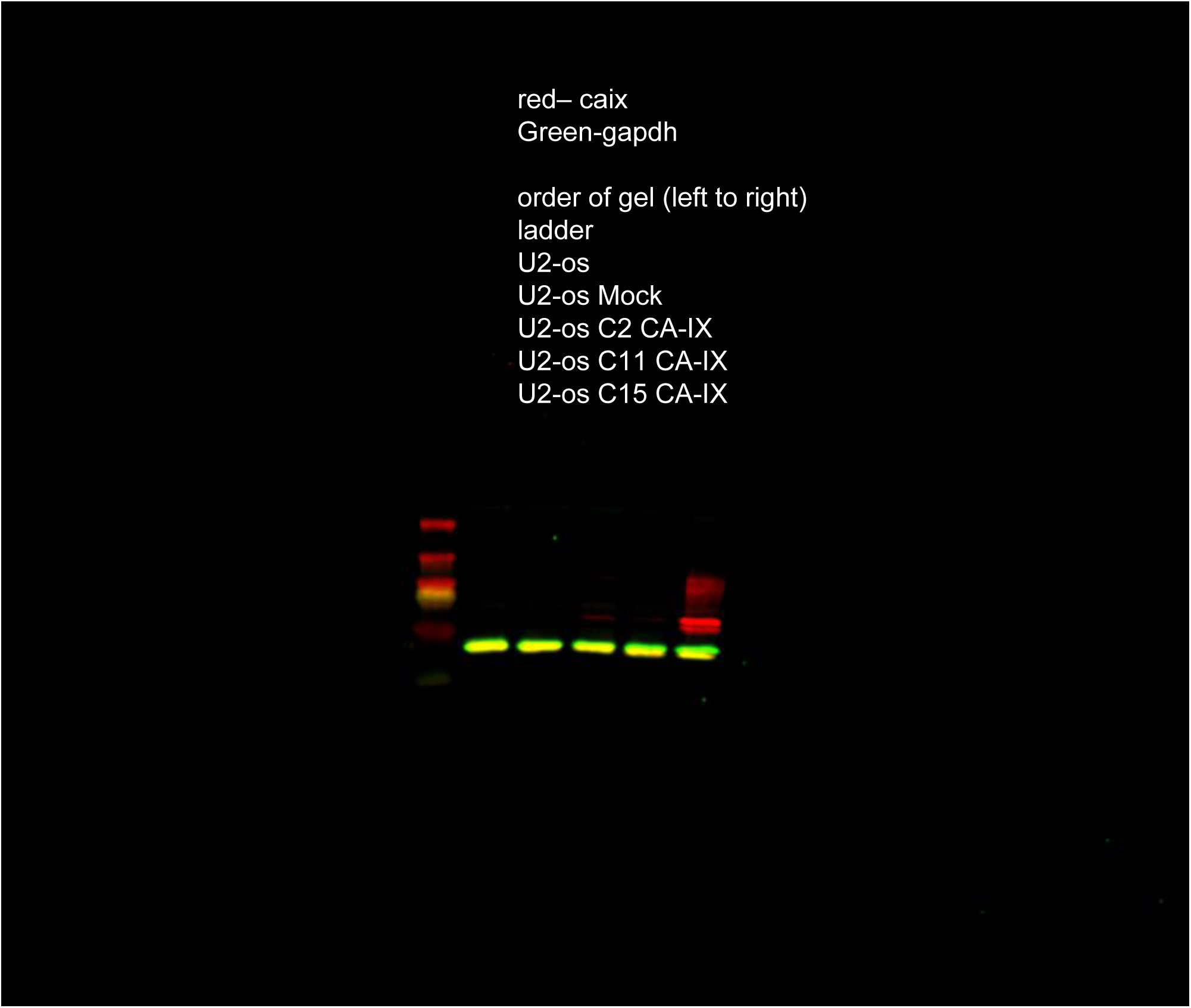

### Raw western – HEK293 CA-IX

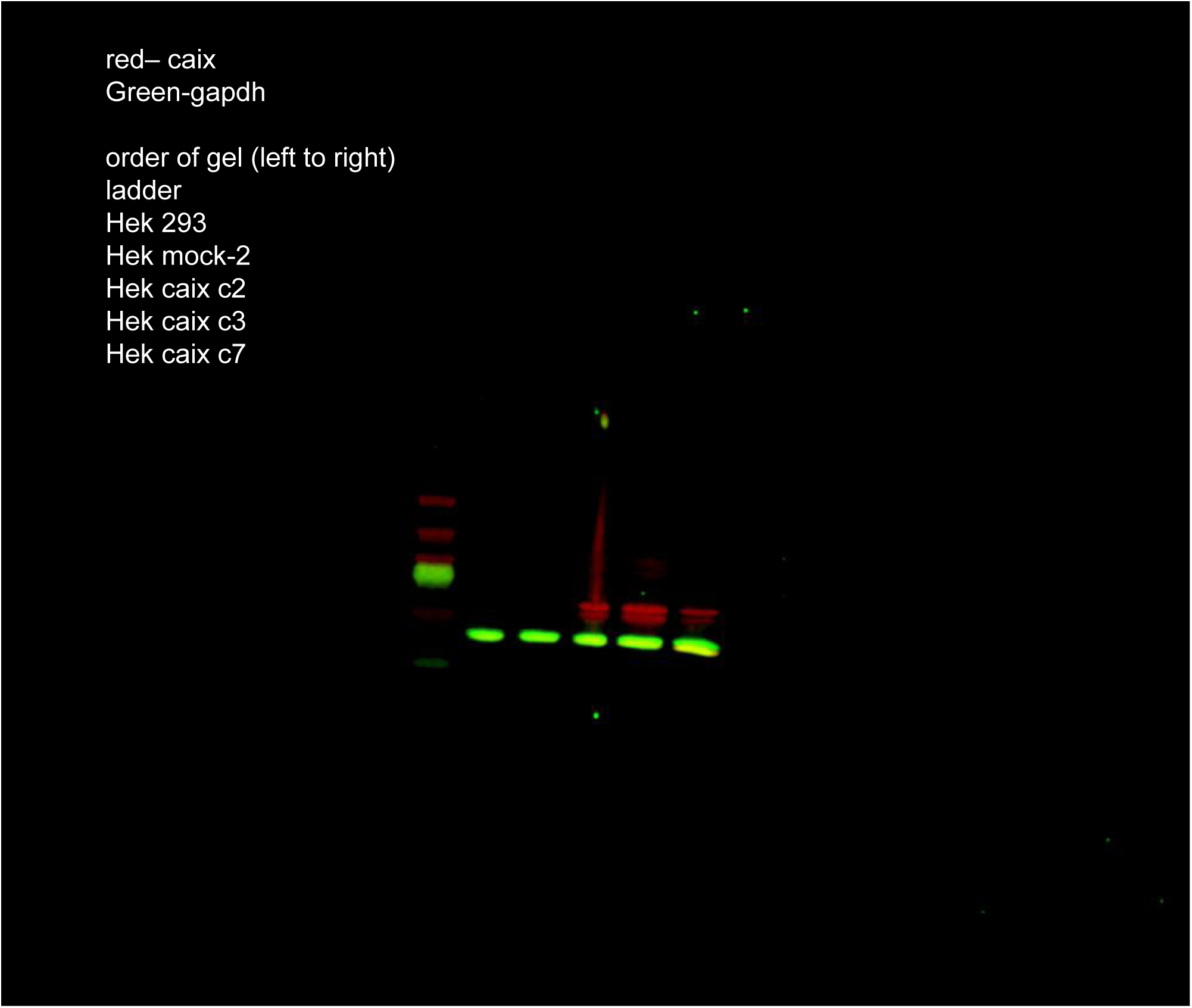

### Raw western – PMA1

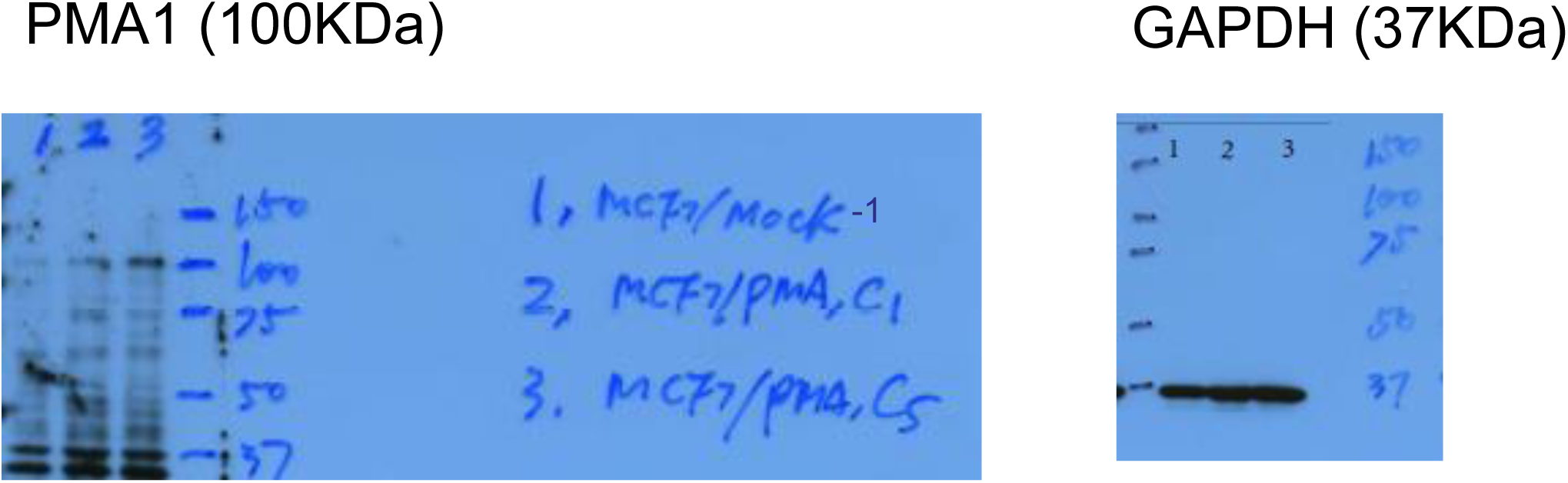

### Matlab code

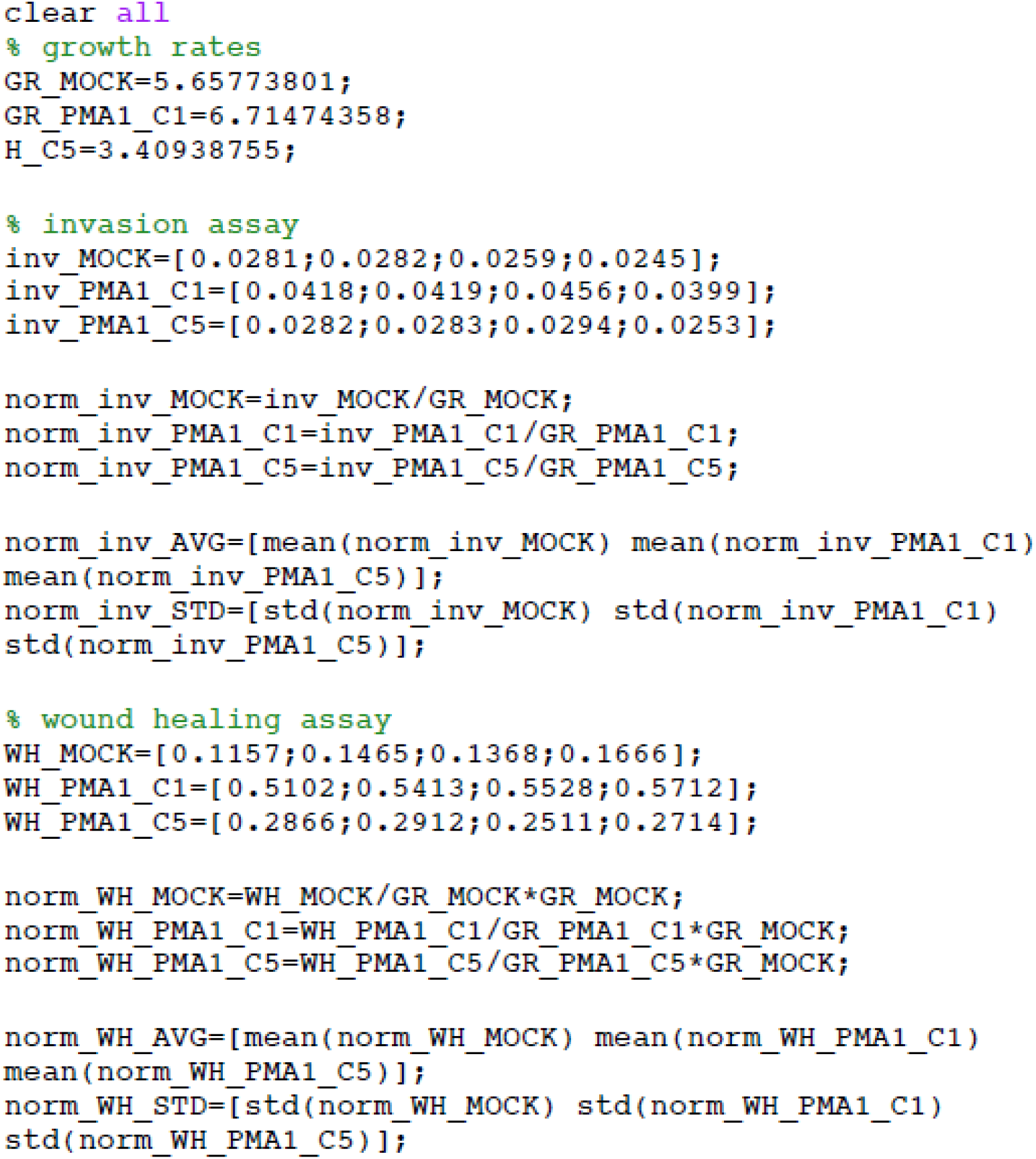

